# Comparative genomics of orobanchaceous species with different parasitic lifestyles reveals the origin and stepwise evolution of plant parasitism

**DOI:** 10.1101/2022.04.13.488246

**Authors:** Yuxing Xu, Jingxiong Zhang, Canrong Ma, Yunting Lei, Guojing Shen, Jianjun Jin, Deren A. R. Eaton, Jianqiang Wu

**Author notes:** Jianqiang Wu, **Email:**. **Author Contributions:** Y.X. and J.W. designed research; J.Z., C.M., and G.S. prepared plant tissues, DNA samples, and RNA samples; Y.X., J.W., Y.L., G.S., and J.J. analyzed data; Y.X., J.W., J.J., and D.E. wrote the paper.

## Abstract

Orobanchaceae is the largest family of parasitic plants, containing autotrophic and parasitic plants with all degrees of parasitism. This makes it by far the best family for studying the origin and evolution of plant parasitism. Here we provide three high-quality genomes of orobanchaceous plants, the autotrophic *Lindenbergia luchunensis* and the holoparasitic plants *Phelipanche aegyptiaca* and *Orobanche cumana*. Phylogenomic analysis of these three genomes together with those previously published and the transcriptomes of other orobanchaceous species, created a robust phylogenetic framework for Orobanchaceae. We found that an ancient whole-genome duplication (WGD; about 73.48 Mya), which occurred earlier than the origin of Orobanchaceae, might have contributed to the emergence of parasitism. However, no WGD events occurred in any lineage of orobanchaceous parasites except for *Striga* after divergence from their autotrophic common ancestor, suggesting that, in contrast to previous speculations, WGD is not associated with the emergence of holoparasitism. We detected evident convergent gene loss in all parasites within Orobanchaceae and between Orobanchaceae and dodder *Cuscuta australis*. The gene families in the orobanchaceous parasites showed a clear pattern of recent gains and expansions. The expanded gene families are enriched in functions related to the development of the haustorium, suggesting that recent gene family expansions may have facilitated the adaptation of orobanchaceous parasites to different hosts. This study illustrates a stepwise pattern in the evolution of parasitism in the orobanchaceous parasites, and will facilitate future studies on parasitism and the control of parasitic plants in agriculture.

## Introduction

Most plants are autotrophs. However, about 4750 angiosperm species are parasitic (Nickrent, 2020), and completely or partly depend on plant hosts to survive, from which they extract water and nutrients. Some parasitic plants still have chlorophyll and photosynthesize; these are known as hemiparasitic plants. Others can no longer photosynthesize, and are known as holoparasites. According to whether hosts are needed for the completion of life cycles, parasitic plants can also be classified as facultative and obligate (Westwood et al., 2010). In angiosperms, parasitic plants independently evolved 12 or 13 times from autotrophic plants (Westwood et al., 2010). In eight parasitic plant lineages, hemiparasitic species became extinct, leaving only holoparasites remaining (Westwood et al., 2010). However, the Orobanchaceae (Lamiales), the largest family of parasitic plants, comprises facultative hemiparasitic, obligate hemi-and obligate holoparasitic species (Supplemental Figure 1) (Westwood et al., 2010; Joel et al., 2013). Orobanchaceae even contains three genera of autotrophs: *Rehmannia, Triaenophora*, and *Lindenbergia*, with *Lindenbergia* being the closest relative of the parasitic lineages (Li et al., 2019). Furthermore, some of the orobanchaceous parasites are economically important, and are devastating pests in agriculture. For example, the annual economic loss caused by the witchweed (*Striga*) and broomrapes (*Orobanche* and *Phelipanche*) is over 10 billion US dollars in Asia and Africa (Twyford, 2018). Orobanchaceae is thus the best extant system in which to study the evolutionary transition to plant parasitism, as well as to develop genomic resources for future agricultural applications (Westwood et al., 2010).

**Figure 1.**
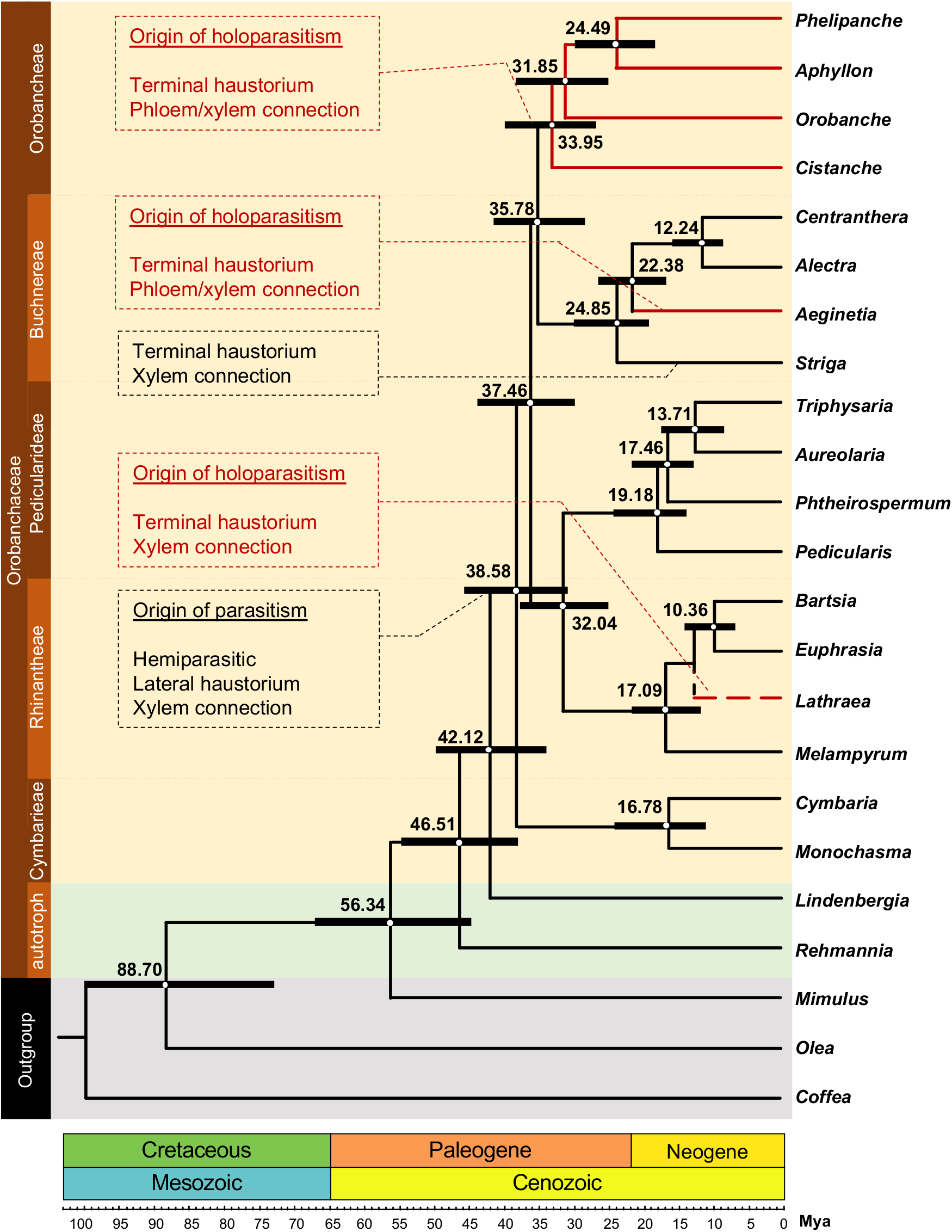
Phylogenetic and molecular clock dating analysis of the major clades in the Orobanchaceae. The phylogenetic tree was constructed using low-copy OGs from 19 orobanchaceous species and three outgroup plants. A gray background indicates autotrophic outgroups; a light grayish-green background represents orobanchaceous autotrophic plant clades; a pale-yellow background represents orobanchaceous parasitic plants; red branches indicate holoparasites. The red dashed line for *Lathraea* highlights that the position of this plant is inferred from the work of Li et al. (2019). The numbers near the internodes represent divergence times (Mya) based on molecular clock dating, and black bars at the internodes represent 95% HPD for divergence times. All internodes are 100% supported. Important evolutionary events are shown in the dashed boxes. The vertical bar on the left indicates five monophyletic clades (Cymbarieae, Orobancheae, Rhinantheae, Buchnereae, and Pedicularideae) and two autotrophic clades (Rehmannieae and Lindenbergieae, which are shown as “autotroph”). The bar on the bottom indicates the geologic time scale.

The phylogenetic relationships among the major lineages of Orobanchaceae remain unclear. Previous research by McNeal et al. (2013) split the family into six major clades based on three nuclear and two plastid markers, while a more recent study by Li et al. (2019) analyzed five additional markers and recognized eight major clades, with incongruent relationships compared to the previous treatment. Following the Angiosperm Phylogeny Group nomenclature, the Li et al. (2019) treatment recognized Rehmannieae, Lindenbergieae, Cymbarieae, Orobancheae, *Brandisia*, Rhinantheae, Buchnereae, and Pedicularideae; furthermore, in Li et al. (2019), the holoparasitic clade Orobancheae was placed as the sister to all other parasitic Orobanchaceae, and the authors proposed that holoparasitism may have evolved early in Orobanchaceae.

The evolution of parasitism requires major morphological and physiological innovations to perceive and recognize potential host plants and to form parasitic connections. Parasitic plants have evolved a unique organ, the haustorium, using which they can attach and penetrate host tissues (Yoshida et al., 2016). The haustoria of many hemiparasites, such as *Triphysaria* and *Striga*, fuse with the xylem of host plants, enabling these hemiparasites to obtain water and nutrients from the host xylem. Some holoparasites fuse their haustorial xylem and phloem with that of their hosts, and symplastic continuity between parasite and host phloem tissues occurs by means of plasmodesmatal connections (Yoshida et al., 2016). The obligate hemiparasites and holoparasites in the Orobanchaceae can perceive strigolactones released from host roots, which activate the germination process in the parasitic plants (Cook et al., 1966; Cook et al., 1972). Recent studies have revealed that in obligate parasites of this family, strigolactones released by the host roots are perceived by the gene *KAI2d*, which appears to have evolved from the karrikin receptor gene *KAI2* by duplication and neofunctionalization. The interaction between the strigolactones and *KAI2d* enables the parasites to detect the existence of host plants in the vicinity (Conn et al., 2015; Tsuchiya et al., 2015; Toh et al., 2015). The evolution and development of haustoria have drawn much attention (Yoshida et al., 2016). Yang et al. (2015) comparatively analyzed the transcriptomes of haustorial tissues from three orobanchaceous species *Triphysaria versicolor, Striga hermonthica*, and *Phelipanche aegyptiaca* and found a core set of parasitism genes, many of which underwent gene duplications or reconfiguration of transcriptional regulation. Previous studies on *Cuscuta australis* (Convolvulaceae) (Sun et al., 2018), which is a shoot parasitic plant, the root parasite *Striga asiatica* (Orobanchaceae) (Yoshida et al., 2019), and an endoparasite *Sapria himalayana* (Rafflesiaceae) (Cai et al., 2021) suggested that gene loss is common in the evolutionary histories of parasitic plants. Wicke et al. (2016) studied the plastomes of various species in the Orobanchaceae, and found that the plastomes of orobanchaceous parasites, particularly those of holoparasites, had clearly experienced large-scale gene loss, leading to most of the photosynthesis-related genes being missing. Gene loss, such as loss of the plastome *ndh* genes, was detected even in the orobanchaceous hemiparasites (Wicke et al., 2016). The degree of gene loss is generally considered to be positively associated with the degree of parasitism (Cai et al., 2021), as a greater degree of parasitism requires higher dependency on the host and fewer physiological processes are needed by the parasite itself. Gene loss may therefore enable the parasite to synchronize its own physiology with that of the host, probably resulting in better fitness (Shen et al., 2020).

To date, genomes from five orobanchaceous species have been released: the autotrophic *Rehmannia glutinosa* (Ma et al., 2021), the obligate hemiparasite *Striga asiatica* (Yoshida et al., 2019), and the facultative hemiparasites *Euphrasia arctica* (Becher et al., 2020), *Phtheirospermum japonicum* (Cui et al., 2020) and *Pedicularis cranolopha* (Jin and Eaton, 2022). However, the genomes of orobanchaceous holoparasites are still lacking. In this study, we sequenced and annotated new reference genomes of the autotroph *Lindenbergia luchensis* and two economically important holoparasitic pests *Orobanche cumana* and *Phelipanche aegyptiaca*. Here we present the largest comparative analysis of genomes of orobanchaceous species to date. A robust phylogeny was reconstructed and the genome evolution of these orobanchaceous species was analyzed in detail, including the histories of genome duplication, gene family expansion, and gene loss, as well as the evolution of haustorium-related genes. Our analyses yielded a new scenario for the origin and evolution of parasitism in Orobanchaceae. This study fills the gap in our understanding of the evolution of plant parasitism and provides novel genomic resources for the most diverse and economically important parasitic plant clade.

## Results

### Sequencing, assembly, and annotation of the genomes of *Lindenbergia luchunensis, Orobanche cumana*, and *Phelipanche aegyptiaca*

Using a Nanopore platform, 36.47Gb data were generated for *Lindenbergia luchunensis* (Llu), and 158.30 Gb and 396.00 Gb data were generated respectively for *Orobanche cumana* (Ocu) and *Phelipanche aegyptiaca* (Pae) on a PacBio Sequel II platform in the CLR mode. Short reads (38.19 Gb, 141.20 Gb, and 329.33 Gb, respectively) were produced for Llu, Ocu, and Pae using an MGI-SEQ 2000. Genome survey analysis using the short reads indicated low degrees of heterozygosity (Supplemental Table 1). Long reads were thus directly used for *de novo* assembly of the genomes, and the short reads were used to correct the genome assemblies, forming contigs. The total length of the Pae genome assembly was 3876.90 Mb (N50 9.97 Mb). Using the Hi-C data, we obtained chromosome-level Llu and Ocu genomes (Llu: 20 pseudochromosomes, 212.21 Mb; Ocu: 19 pseudochromosomes, 1417.94 Mb, Supplemental Table 1). The accuracy of each of the three genomes was found to exceed 99.9% (see Supplemental Methods and Supplemental Table 1). We then annotated repetitive sequences, including simple repeats, transposable elements, and RNA genes. In the genomes of Llu, Ocu, and Pae, the repetitive sequences occupied 30.72%, 79.82%, and 84.40%, respectively (Supplemental Table 2). Mapping the RNA-seq data (Supplemental Table 3) from different organs to the respective genomes resulted in 29,669, 42,525, and 50,484 protein-coding genes in Llu, Ocu, and Pae, respectively. Benchmarking universal single-copy orthologs (BUSCO) analysis indicated that the completeness of these three genomes was 97.1% (Llu), 80.5% (Ocu), and 76.3% (Pae) (Supplemental Table 4). The low completeness values of the genes in Ocu and Pae genome is due to the large-scale gene losses seen in Ocu and Pae (see below for analysis), which is associated with their holoparasitic lifestyle.

### Phylogenetic relationships in Orobanchaceae based on nuclear gene sequences

Previous phylogenetic analyses suggested that there are eight major clades in the Orobanchaceae, but that the phylogenetic relationships among the clades were still uncertain (Schneeweiss, 2013; McNeal et al., 2013; Li et al., 2019). The nuclear genes from the 19 available genomes/transcriptomes of Orobanchaceae species (Supplemental Table 5), which cover seven of the eight clades of Orobanchaceae, and three outgroup species (Supplemental Table 5) were used in our phylogenetic analysis (see Supplemental Methods). Finally, after filtering out those orthogroups (OGs) which had low quality and/or high copy numbers, 907 OGs were selected for phylogenetic reconstruction. Coalescent and concatenation methods yielded identical tree topologies, with 100% support for all the nodes (Figure 1). In this phylogenetic tree, which is consistent with previous findings (Wolfe et al., 2005; Schneeweiss, 2013; McNeal et al., 2013; Li et al., 2019), the autotrophic lineage Rehmannieae is sister to all the other orobanchaceous lineages, and another autotrophic lineage Lindenbergieae is the sister clade of all the parasitic plant lineages (Figure 1). Within the group of parasitic plants, Cymbarieae is the sister group to all the remaining lineages, which differs from the results of Li et al. (2019), which was based only on a few nuclear genes. Buchnereae and Orobancheae are sister clades, as are Pedicularideae and Rhinantheae (Figure 1). The topology of these four clades is distinct from the results of all previous studies (Wolfe et al., 2005; Schneeweiss, 2013; McNeal et al., 2013; Li et al., 2019), although it is similar to the topologies based on the *matK* and *rps2* genes (McNeal et al., 2013).

Several previous studies indicated that parasitic plants have higher nucleotide substitution rates than typical autotrophic plants (Lemaire et al., 2011; Bromham et al., 2013); however, in our phylogenetic tree, only the branch length of the holoparasitic plant *Aeginetia indica* in the clade Buchnereae is exceptionally long, while the branches of all holoparasitic plants in the clade *Orobancheae* were similar to those of the autotrophic plants (Supplemental Figure 2). We calculated the K_s_ values of all the lineages in Orobanchaceae, and indeed no branches showed high substitution rates (Supplemental Figure 3).

Based on this tree, we estimated the divergence times for all the nodes with a Bayesian molecular dating analysis. The estimated origin of the Orobanchaceae is about 46.51 million years ago (Mya; 95% highest posterior density (HPD), 37.91-55.39 Mya), and the emergence of parasitism predates 38.58 Mya (95% HPD, 31.27-45.81 Mya). Our analysis also indicated that holoparasites evolved independently at least three times in the Orobanchaceae (Figure 1), a result similar to that of Fu et al. (2017), and of these three lineages of holoparasites, Orobancheae appeared the earliest, at least 33.95 Mya (95% HPD, 27.41-40.43 Mya).

### Genome evolution in the Orobanchaceae

Genome sizes and chromosome numbers vary in the Orobanchaceae, and they seem to be positively associated with the degrees of parasitism (Joel et al., 2013; Lyko and Wicke, 2021). It is plausible that multiple rounds of WGD events occurred during the evolution of orobanchaceous species (Joel et al., 2013; Lyko and Wicke, 2021). To comprehensively reveal the WGD events in the Orobanchaceae, six species with reference genomes (Pae, Ocu, and Llu from this study; *Striga asiatica* (Sas), *Phtheirospermum japonicum* (Pja), and *Pedicularis cranolopha* (Pcr) from published/released data) and four outgroup species (*Mimulus guttatus* (Mgu; Lamiales), *Solanum lycopersicum* (Solanales, lamiids), *Coffea canephora* (Gentianales, lamiids), and *Arabidopsis thaliana* (Brassicales, rosids)) were chosen for colinear fragment comparisons. Instead of the commonly used methods based on colinear fragments obtained by homology searches, an analysis based on “orthologous colinear fragments (OCFs)” was specifically designed for this study (see Supplemental Methods and Supplemental Figure 4), which enabled us to investigate all possible WGD events during the evolution of the Orobanchaceae.

Our analysis based on OCFs from all species pairs (Figure 2) indicated that the Sas lineage experienced an independent WGD event (named *α*^*B*^ event; Figure 2), which was consistent with a previous study (Wickett et al., 2011; Yoshida et al., 2019). However, no WGD events were detected in any other orobanchaceous lineages, including those containing Pae and Ocu (Figure 2). The ancestor of Orobanchaceae did not experience any independent WGD events, even though there was a WGD event during the evolution of the common ancestor of Orobanchaceae and *Mimulus*, which is named the *β*^*L*^ event (Figure 2). Notably, the detected WGD events in the outgroup species are consistent with previous reports (Simillion et al., 2002; Sato et al., 2012; Denoeud et al., 2014), supporting the reliability of our novel WGD analysis pipeline. Next, we selected four nodes (S1-S4 in Figure 2) within the Lamiales clade, and all the orthologous genes originating from these speciation events were selected for calculation of the K_s_ values (Supplemental Figure 5). We found that the peak values of the K_s_ distributions had a linear relationship with the divergence times of these nodes (*R*^2^ = 0.9926, Figure 2). Therefore, the K_s_ values from the paralogous genes resulting from the *α*^*B*^ and *β*^*L*^ events were used to infer the times of these two events: *α*^*B*^ and *β*^*L*^ occurred at 32.64 ± 6.58 and 73.48 ± 6.58 Mya, respectively (Figure 2). The *α*^*B*^ event likely occurred in the Buchnereae soon after it diverged from the Orobancheae at 35.78 Mya (95% HPD, 28.97-42.52 Mya; Figures 1 and 2). The occurrence of the *β*^*L*^ event was close to the Cretaceous-Paleocene boundary, and the mass extinction event at this time (Schulte et al., 2010) when many plant lineages experienced WGD events (Fawcett et al., 2009; Wu et al., 2020). We speculated that the *β*^*L*^ event may have played an important role in enabling the common ancestor of Mgu and the Orobanchaceae to survive the extreme environmental changes at the Cretaceous-Paleocene boundary.

**Figure 2.**
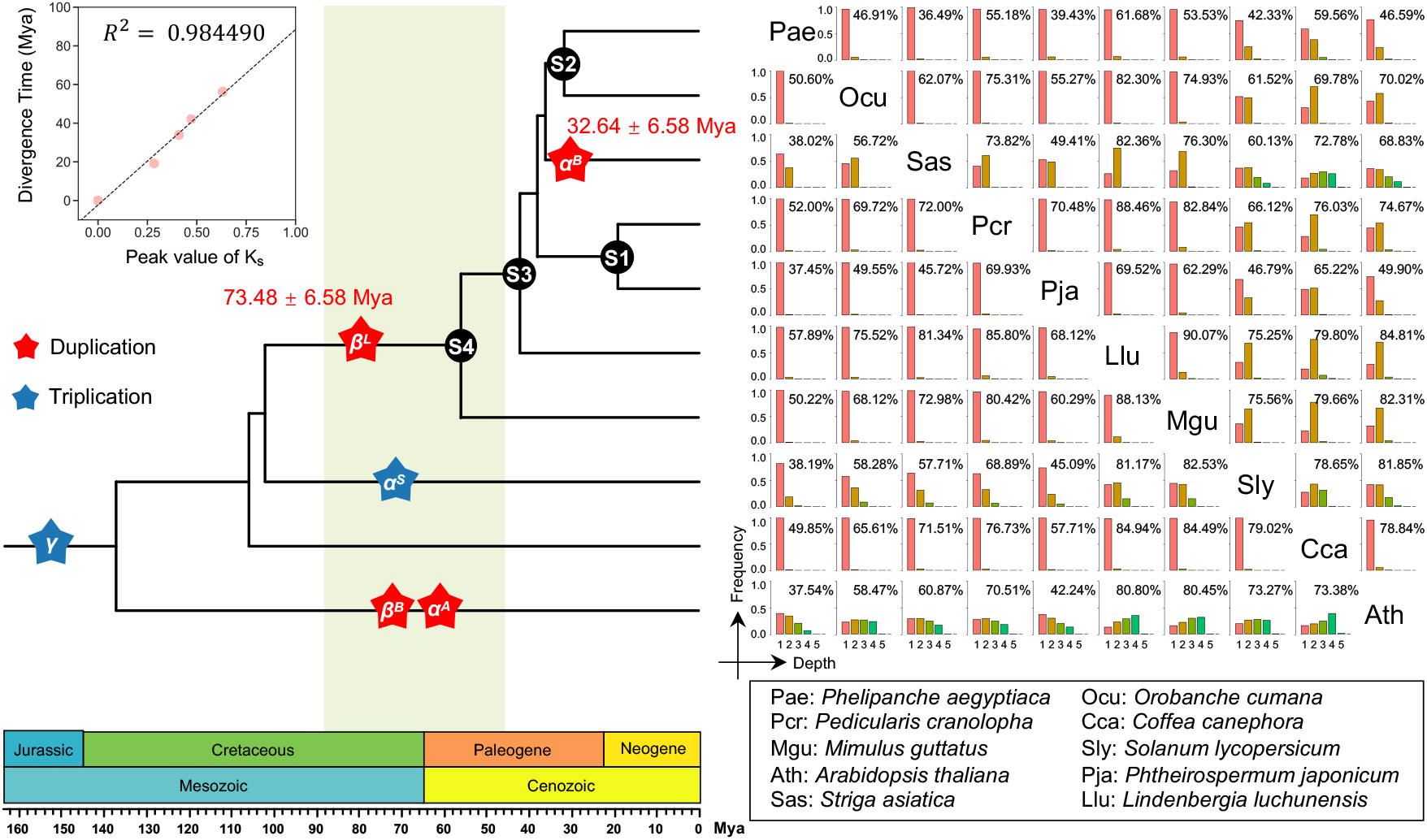
Genome evolution in the Orobanchaceae. The palaeopolyploidization in the Orobanchaceae and outgroup lineages. The right histogram matrix indicates the coverage depths of orthologous colinear fragments (OCFs) from pairwise species comparisons. The OCFs from a given species pair were mapped respectively to each species to create two histograms indicating the coverage depths of OCFs (X-axis: coverage depths of OCFs, Y-axis: relative frequency). The max depths of OCFs mapped to species B can be used to infer the WGD events in species A after speciation of species A and B, and vice versa (see Supplemental Methods). In the matrix, each row and column of the histogram indicate the two species used to create the OCFs, and the species of the column is the one that was the mapping-target species. The percentage in the upper right corner of each histogram gives the proportion of genes covered by OCF in the target species to the total genes. The WGD events inferred from the histogram matrix are marked with stars in the species tree on the left (blue: triplication, red: duplication), and the positions of the stars represent the times of the WGD events, which were inferred from the peak value of K_s_. The black circles on the species tree represent the four speciation events which were used to calculate the correlation between K_s_ values and divergence times (see Supplemental Methods). The correlation between the peak value of K_s_ and divergence times is shown in the upper left corner.

Knowledge of chromosome numbers is important for understanding the genome evolution of eukaryotes. The chromosome numbers in the Orobanchaceae vary (Joel et al., 2013; Lyko and Wicke, 2021). Based on the chromosome-level genome data from Ocu, Pcr, Llu and the outgroup species Mgu, we obtained 18 protochromosomes by reconstructing the ancestral karyotype of the Orobanchaceae (Supplemental Figure 6). Analysis of the syntenic chromosomal fragments indicated that Llu and Ocu exhibited good collinearity with the ancestral protochromosomes (Supplemental Figures 6 and 7). For example, Chromosome 9 of Ocu and Chromosome 5 of Llu appeared to be very similar to the ancestral Chromosome 8. However, Pcr has only eight chromosomes and its chromosomes are likely to have experienced at least 11 fusion events.

Orobanchaceous species have variable genome sizes, ranging from 223 Mb to 10.7 Gb, (Bai et al., 2012; Joel et al., 2013; Lyko and Wicke, 2021). Of the six orobanchaceous species we studied (Llu, Pja, Pcr, Sas, Ocu, and Pae), Llu has the smallest genome (223 Mb), while Pae has the largest (3992 Mb), 20 times bigger than the Llu genome. Even though recent WGD can rapidly increase genome size, our analysis ruled out recent WGD events in the evolution of these six species. Thus, to gain insight into the driving force behind the genome expansions in these parasites, we compared the genome sizes of Llu, Pja, Pcr, Sas, Ocu, and Pae and the repetitive elements (REs) in their genomes (Figure 3A). The large genome sizes mainly resulted from increased numbers of REs. In the genomes of Ocu, Pja, and Pcr, which are about 1 Gb, 64-79% of the genomes are made up of REs; Pae has the largest genome (3.877 Gb), and 84% of the Pae genome comprises REs, with the Long Terminal Repeats (LTRs) family being the most abundant. Notably, analysis of the sequences of the LTRs in Llu, Pcr, Ocu, and Pae suggested that the subfamilies of LTRs expanded are species-dependent, especially during the very recent extreme extents in Pae and Ocu (Supplemental Figure 8). To determine whether the massively increased numbers of LTRs could affect the coding regions in these species, we compared the intron sizes in Pae and Llu, which have the largest and smallest genomes, respectively, of the orobanchaceous species included in this study. We found that 73.23% of intron length variations were less than 1-fold, regardless of genome sizes (Figure 3B). Moreover, the RE contents in the gene flanking regions became larger as genome size increased (Figure 3C). Dramatic changes in the flanking sequences of a large number of genes may have caused genome-wide changes in gene expression. We also noticed that even though no WGD events occurred, Pja, Pcr, Pae, and Ocu, which have very large genomes, had many more genes than does Llu, and that this was mainly due to tandem or segmental gene duplication (30.96%-56.33%; Supplemental Table 6), but not LTR-assisted “copy-paste” duplication (6.55%-17.40%; Supplemental Table 6). Because Chromosome 9 of Llu and Chromosome 5 of Ocu are homologous, we specifically inspected the differences between these two chromosomes. Ocu’s Chromosome 5 is eight times larger than is Chromosome 9 of Llu (82 Mb vs 10 Mb). We found many Class I retrotransposons, including LTRs, in the proximal region of Chromosome 5 of Ocu, where gene densities are relatively low, supporting our hypothesis that expansion of LTRs is the main driving force behind the increased genome sizes in orobanchaceous parasites (Figure 3D). This finding is consistent with a previous conclusion obtained by genome skimming (Piednoël et al., 2012).

**Figure 3.**
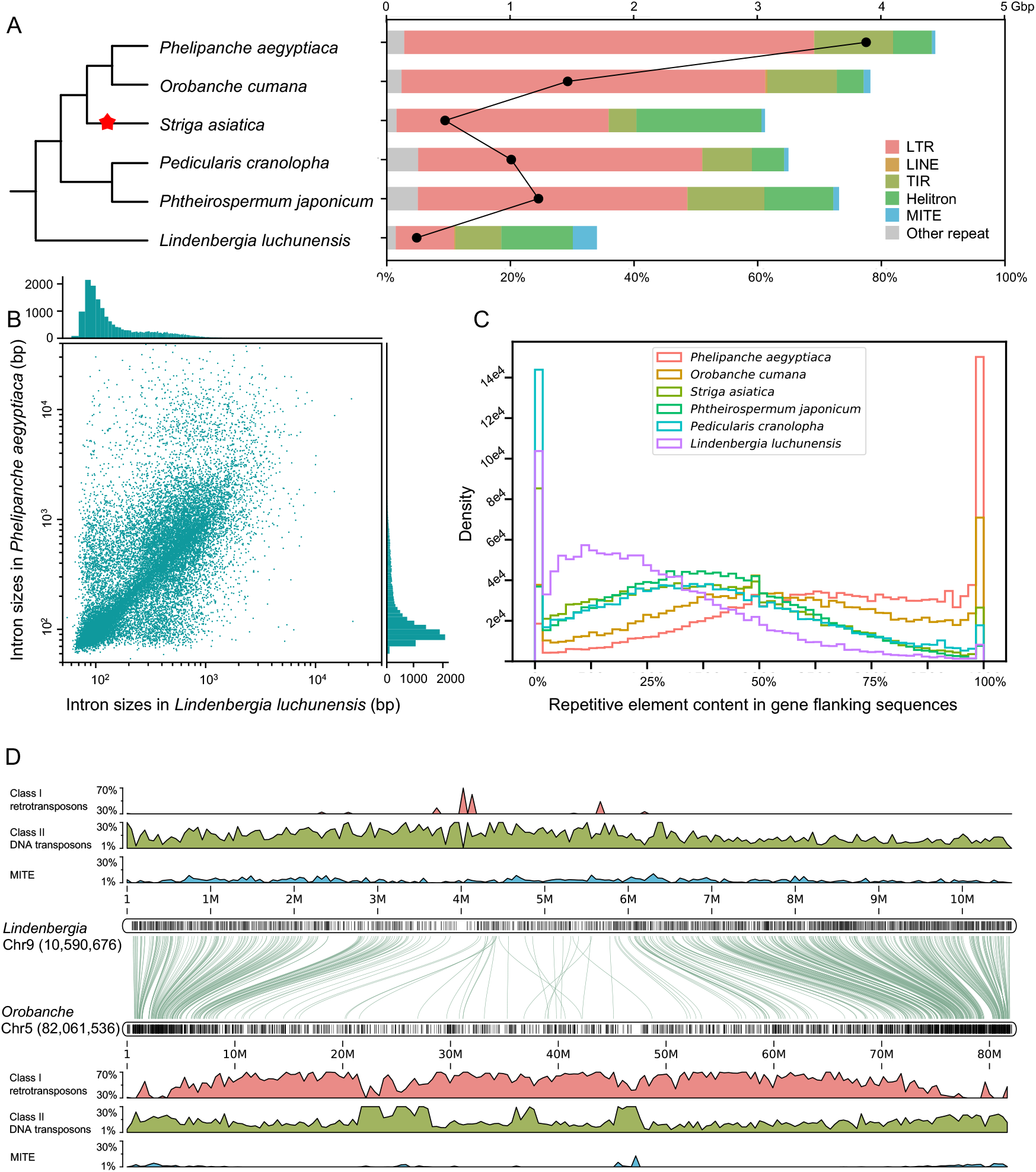
Expansion of repetitive elements in the genomes of species in the Orobanchaceae. (A) Genome sizes and proportions of repetitive elements in six species of Orobanchaceae. The black dots and upper X-axis represent the genome sizes, and the bars and lower X-axis represent the percentages of different types of repetitive elements in the genomes. The WGD event is marked with a red star in the species tree. (B) Size distribution of introns that are conserved in Llu and Pae. Each dot represents an intron, and the histograms on the top and to the right show the size distribution of introns in Llu and Pae, respectively. (C) Repetitive element content in gene flanking regions in six species of Orobanchaceae. (D) Comparison of chromosomal repetitive elements and gene positions in chromosome 9 of *Lindenbergia luchunensis* and chromosome 5 of *Orobanche cumana*. Black lines in the boxes indicate the positions of coding genes on the chromosomes; area graphs indicate the proportion of different types of repeat sequences in the corresponding chromosomal regions; curved lines connect orthologous gene pairs. The length of the bin used for plotting is 1/1000 of the total chromosome length.

### Evolution of genes and gene families in the Orobanchaceae

We analyzed all the gene families in the orobanchaceous species (Pae, Ocu, Sas, Pja, Pcr, and Llu), *Cuscuta australis* (Cau), and their relatives (for orobanchaceous species: Mgu and *Sesamum indicum*; for Cau: *Ipomoea nil* and *Solanum lycopersicum*), as well as two outgroup species, *Coffea canephora* and *Arabidopsis thaliana* (Dataset S1). We found that, like Cau, all parasites in the Orobanchaceae experienced frequent gene family contractions; however, many gene families in orobanchaceous parasites exhibited severe expansions, which are rare in Cau (Figure 4). Based on the phylogenetic tree (Figure 1), we calculated the numbers of gene families that underwent significant expansions and contractions across all branches (Figure 4). Contractions were frequent when parasitism and obligate parasitism first appeared in the Orobanchaceae, and thereafter, most lineages appeared to have undergone independent gene family contractions. Most of the gene family expansions occurred independently in different species. Of the expanded gene families, genes related to transcriptional regulation, transport, cell wall formation, phloem/xylem histogenesis, and lateral root formation were enriched (Dataset S2), and these genes may be involved in the evolution of the haustorium.

**Figure 4.**
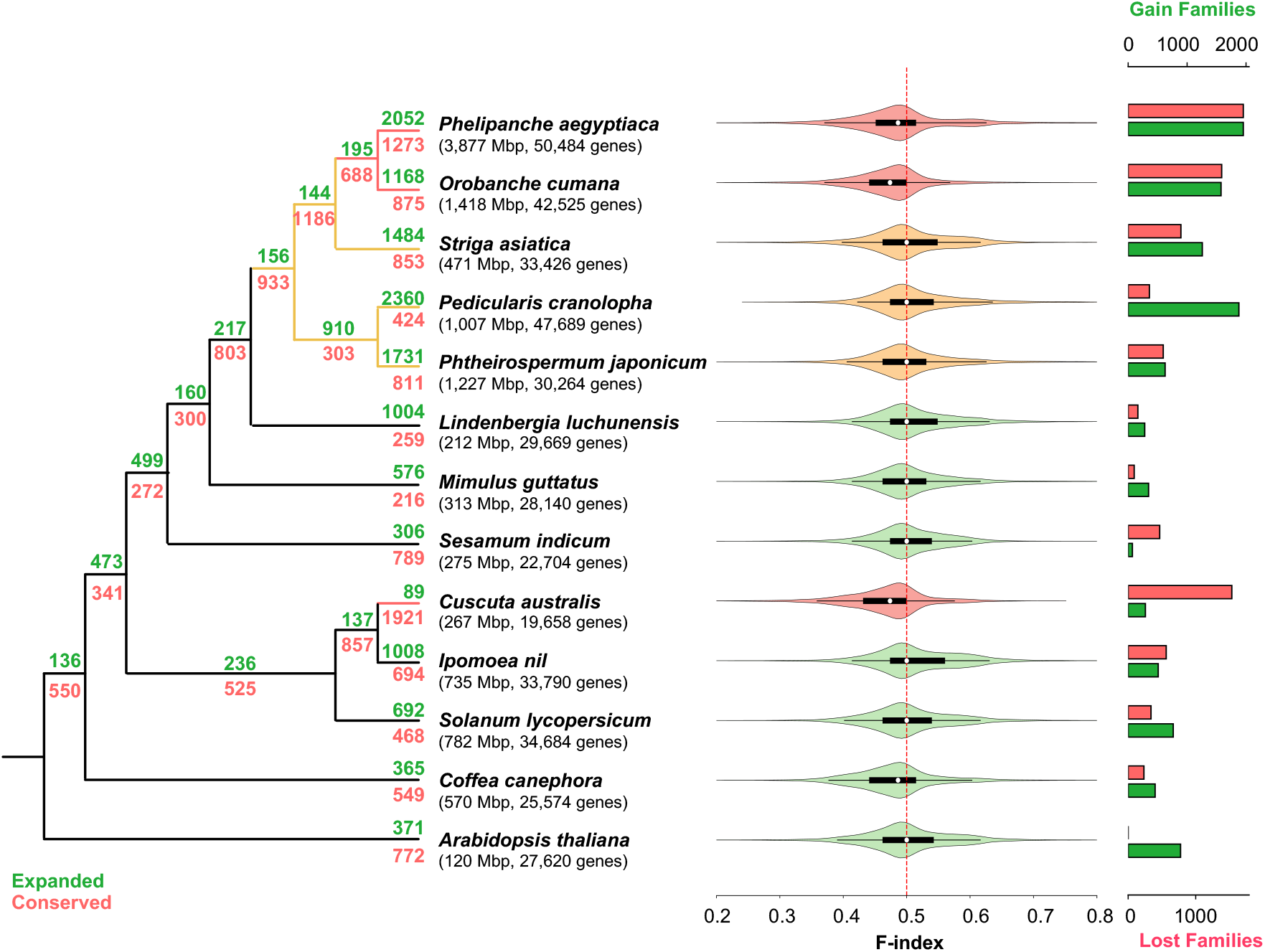
Gene family expansion and contraction in the Orobanchaceae. On each branch of the species tree, the green number indicates the numbers of expanded gene families, and the red number indicates the number of conserved gene families. Each density curve in the violin plot shows the F-index (see Supplemental Methods) distribution of conserved gene families in each of the species indicated to the left. In the density curves, soft red, soft orange, and grayish green indicate holoparasitic, hemiparasitic, and autotrophic plant species, respectively. The bar plot on the right shows the numbers of conserved gene families which were completely lost in each species (red bars with lower X-axis) and the numbers of species-specific gene families (green bars with upper X-axis).

Furthermore, we screened the transcriptomic data (Supplemental Table 7) from Pja, Sas, and Pae for genes with high expression levels in the haustoria (where expression level of a given gene was at least 1-fold greater in haustorium tissues than in other tissues, see Supplemental Methods), and found that at least 40.92% of these haustorially highly expressed genes (HHEGs) evolved through duplication after the emergence of the most recent common ancestor (MRCA) of Cca and the Orobanchaceae (Figure 5A, Supplemental Table 8). Next, these duplicated genes were used to place the duplication events in the phylogeny (Figure 5A, Supplemental Table 9): 28.03%-36.34% of the duplication events occurred in the MRCA of Mgu and the Orobanchaceae (S4 in Figure 2), which speciated after the *β*^L^ WGD event (Figure 2). To quantify how many of the duplication genes among the HHEGs resulted from the *β*^L^ WGD event, we extracted the colinear fragments where the gene duplicates were located and found that 96.91% (439/453) of these duplicated genes were located in colinear fragments caused by the *β*^L^ event. Thus, at least 11% of the HHEGs originated from the *β*^L^ event. Most of the other duplication events occurred during the specific evolution of Pja, Sas, and Pae, but not in the common ancestor of the orobanchaceous parasites (Figure 5A, Supplemental Table 9). Therefore, the *β*^L^ event was associated with the origin of many haustorium-related genes.

**Figure 5.**
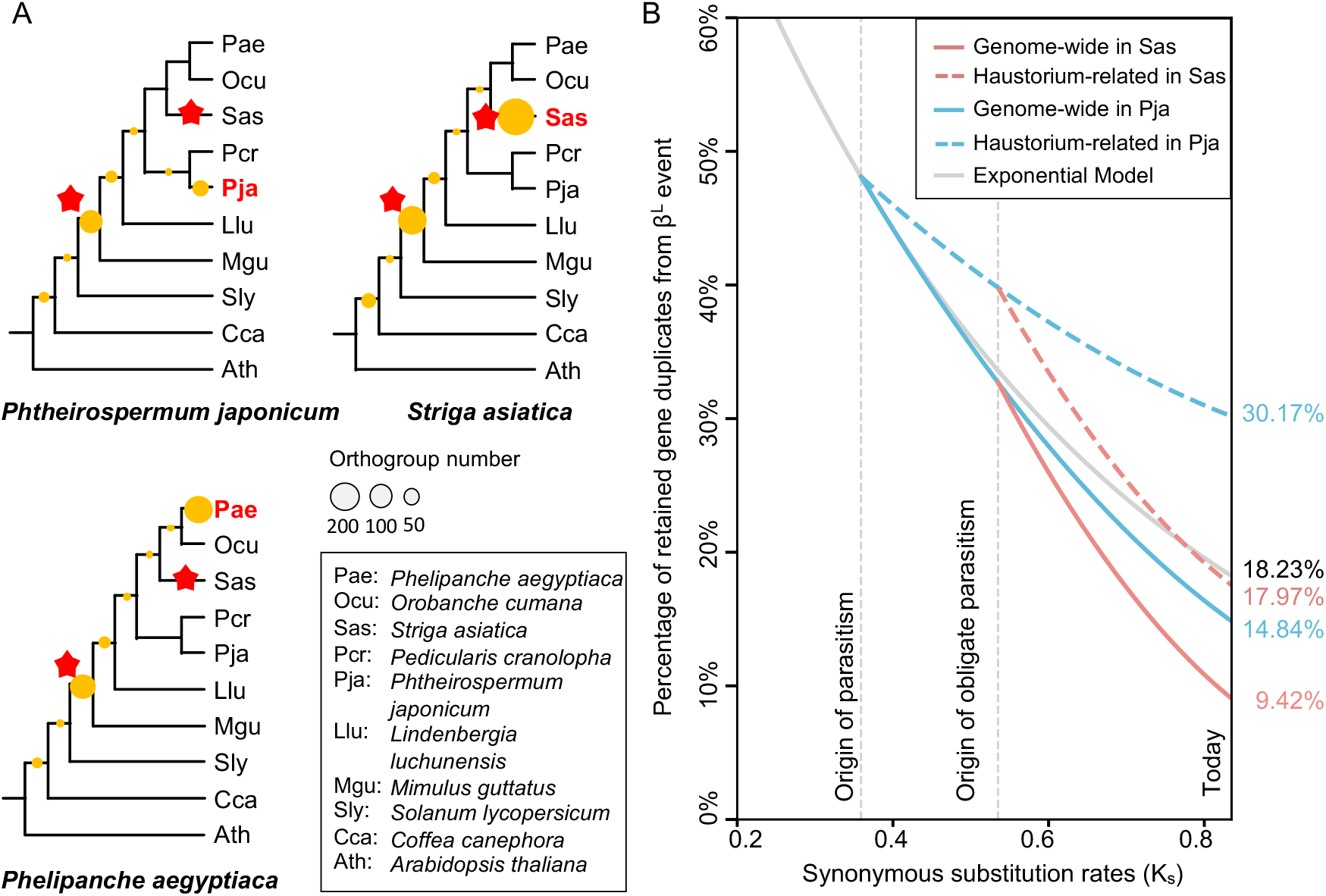
The role of the ancient WGD event *β*^*L*^ in the emergence and evolution of parasitism in the Orobanchaceae. (A) Phylogenetic placement of gene duplications detected in the highly expressed genes in the haustorium. The highly expressed genes in the haustorium of each species were identified. Gene trees were constructed for all the gene families comprising these highly expressed genes. The duplication events were inferred from the topologies of the gene trees. The size of the yellow circle on each tree node represents the number of gene duplication events occurred during that time. The WGD events are marked with red stars in the species tree. (B) Loss of gene duplicates after the *β*^L^ WGD event in *Striga asiatica* and *Phtheirospermum japonicum*. The X-axis (K_s_) represents time after the *β*^L^ WGD event (see Supplemental Figure 5 for details of calculation). The gray curve represents the predicted values based on an exponential model under neutral selection (Supplemental Methods). Red and blue curves were inferred from the predicted initial values and the observed end values respectively in *Striga asiatica* and *Phtheirospermum japonicum* (Supplemental Table 10).

The *β*^L^ event occurred at 73.48 ± 6.58 Mya, while the origin of parasitism was estimated to be at least 38.58 Mya (95% HPD, 31.27-45.81 Mya), ∼ 35 million years after the *β*^L^ event (Figure 2). After a WGD event, the duplicated genes are often rapidly lost (Maere et al., 2005). We used a mathematical model in which the duplicated genes are exponentially lost during evolution under neutral selection (Ren et al., 2018) (see Supplemental Methods) to estimate the fates of duplicated genes resulting from the *β*^L^ event, and then we compared the predicted values with the retained duplicated genes from the *β*^L^ event in Mgu, Llu, Pja, Sas, and Pae. The model predicted that 48.10% of the duplicate genes from the *β*^L^ event would be retained when parasitism emerged, and assuming neutral evolution, that about 18.23% of the duplicated genes should be retained in the extant genomes, a proportion similar to those observed in the autotrophic Mgu and Llu (Figure 5B, Supplemental Table 10). In the genome of the facultative hemiparasite Pja, the retention rate of *β*^L^-generated gene duplicates was 17.97%, a value similar to the value predicted under neutral selection (18.23%). However, when we focused on the HHEGs, the retention rate increased to 30.17% (Figure 5B, Supplemental Table 10), suggesting strong positive selection. In terms of the obligate hemiparasite Sas and holoparasite Pae, only 9.42% and 9.06%, respectively, of the duplicates resulting from the *β*^L^ event were retained (Figure 5B, Supplemental Table 10); these retention rates are much lower than the model predictions, and this might be related to the large-scale gene loss in obligate parasites (see below). Similarly, the retention rates of duplicated HHEGs resulting from the *β*^L^ event decreased to 17.97% and 17.54%, respectively (Figure 5B, Supplemental Table 10). Thus, in both facultative and obligate parasites, the retention rates of gene duplicates from the *β*^L^ event are higher in HHEGs than in genome-wide gene duplicates, suggesting that the *β*^L^ event was important for the origin of the haustorium and parasitism. Furthermore, more of the gene duplicates resulting from the *β*^L^ event were retained in the facultative than in the obligate parasites. It is possible that during the evolution of obligate parasites from their facultative ancestor, some of the gene duplicates were no longer required and were therefore deleted from the genome.

Gene loss is common in parasitic plants (Sun et al., 2018; Cai et al., 2021). We split all the gene families into OGs, and these OGs were thereafter used for gene loss analysis. Gene family contractions were not additionally analyzed in other ways, as comparison of gene loss provides detailed information on the biological significance of the missing genes. A bioinformatic pipeline was also employed to screen for false positives generated by errors of gene annotation and orthogrouping (see Supplemental Methods). As a comparison, Cau was included in the gene loss analysis with five parasitic Orobanchaceae species, as *Cuscuta* and Orobanchaceae are both Lamiids. *Sapria himalayana* (Rafflesiaceae) (Cai et al., 2021) was not included as it is phylogenetically very distant from Lamiids and is an endoparasite. We found that the severity of gene loss was positively associated with the degree of parasitism: the facultative hemiparasites Pja and Pcr lost 2.41% and 2.95%, respectively, of their conserved genes, the obligate hemiparasite Sas lost 5.71%, and the obligate holoparasites Ocu, Pae, and Cau lost 13.11%, 13.85%, and 14.88%, respectively (Figure 6A and B). The history of gene loss events in orobanchaceous parasites was then estimated based on the principle of maximum parsimony, and the common ancestor of all orobanchaceous parasites was found to have lost only 74 genes, while 191 and 777 genes were lost, respectively, during the evolution of the ancestors of obligate parasites and holoparasites (Figure 6A). Notably, even though hundreds of genes were lost during the species-specific evolution of these parasites (Figure 6A), many were convergent (Figure 6A). By comparing the lost genes in the Pedicularideae (Pja and Pcr) and the Orobancheae (Ocu and Pae), we found that 74-122 genes were convergently lost, which was well supported by the hypergeometric test (Figure 6A). Furthermore, in the closely related holoparasites Ocu and Pae, 33.6% and 29.9% of the lost genes were species-specific, and selection analysis indicated that significant portions of these Ocu-and Pae-retained genes, which had been lost in the other species, were under relaxed selection (Supplemental Figure 9). It is possible that Ocu and Pae are still experiencing genes loss, and these genes will be convergently lost in the future due to relaxed selection.

**Figure 6.**
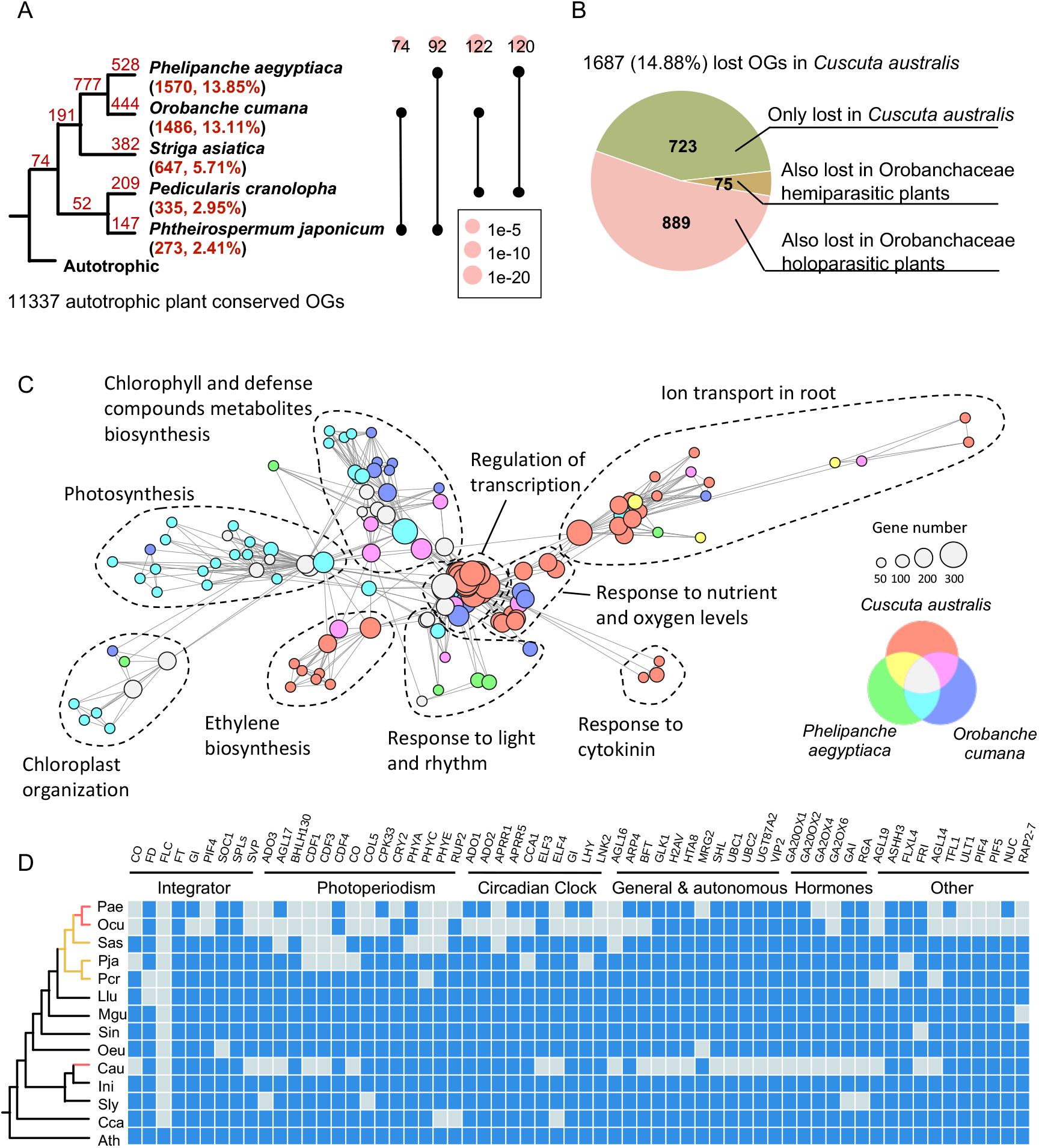
Gene loss in the orobanchaceous parasitic plants and *Cuscuta australis*. (A) History and convergent evolution of gene loss in orobanchaceous parasitic plants. Left: the red numbers on the branches of the species tree represent the numbers of lost OGs estimated based on the principle of maximum parsimony; the number and percentage in brackets under the species name indicate the total number of lost OGs and the proportion of total number of lost OGs to the number of conserved OGs in autotrophic plants, respectively. Right: the numbers on top indicate the numbers of the same lost OGs between the two species indicated by the dumbbells after their divergence from the MRCA; the size of the soft red circle on each number indicates the *p*-value of hypergeometric test. (B) The numbers of commonly and specifically lost OGs in *Cuscuta australis* and orobanchaceous hemiparasitic and holoparasitic plants. (C) Gene ontology (GO) enrichment analysis of lost OGs in *Cuscuta australis, Phelipanche aegyptiaca*, and *Orobanche cumana*. Each node in the networks plot represents the GO term of a biological process that was enriched (p-value < 0.02), and its size indicates the number of OGs associated with the GO term. Nodes were assigned colors depending on whether the gene loss occurred specifically or in common with other species; each edge represents the similarity between two GO terms based on the number of common OGs. (D) Loss of genes related to flowering time regulation in orobanchaceous parasitic plants and *Cuscuta australis*. Grey and blue boxes represent absence and presence of the specific gene, respectively. Pathway information was retrieved from FIOR-ID (http://www.phytosystems.ulg.ac.be/florid/).

We next compared the lost genes in the orobanchaceous parasites and Cau. Among the 1687 lost genes in Cau, more than half (889 genes, 57.14%) were also lost in Pae and Ocu, and 75 genes were also lost in the hemiparasites Sas, Pja, and Pcr (Figure 6B). It is evident that all orobanchaceous parasites and Cau convergently lost various genes, suggesting that some of these gene loss events could contribute to their adaptation to the parasitic lifestyle. To gain further insight into the functions of the lost genes in Cau, Pae, and Ocu, which are all holoparasites, we first performed gene ontology (GO) enrichment analysis on all their lost genes, and the GO terms were then used for interspecies pairwise comparisons. The resulting common and unique GO terms were further categorized using a network analysis (Figure 6C). Pae and Ocu-lost genes exhibited very similar GO terms, while the enriched GO terms from the Cau lost genes appeared to be quite different from those of Pae or Ocu (Supplemental Figure 10). In Cau, most of the GO terms for the lost genes were related to ion transport in roots, ethylene biosynthesis, and response to nutrient and oxygen levels, which is consistent with the rootless morphology of Cau (Figure 6C). The majority of the lost genes in Pae and Ocu were involved in photosynthesis, chloroplast organization, and chlorophyll biosynthesis. Although Cau also lost genes related to these processes, the numbers of GO terms under these processes are markedly fewer than those from Pae or Ocu (Figure 6C).

In the leaves of autotrophic plants, a complex flowering regulatory network perceives endogenous and environmental cues and determines the flowering time of the plant accordingly (Blümel et al., 2015). Given that parasitic plants need to interact with their hosts, and that holoparasites no longer have leaves, which are the organs that determine flowering (Jaeger et al., 2013), parasitic plants are likely to have evolved unique mechanisms to regulate flowering. Genome sequencing of the dodder Cau and a fully mycoheterotrophic orchid *Gastrodia elata* indicated that many flowering-related genes were lost in both species (Sun et al., 2018; Xu et al., 2021). We therefore inspected the genes responsible for the regulation of flowering in all the orobanchaceous parasites included in this study, with Cau included for comparisons (Figure 6D). We found that many important flowering regulatory genes were indeed missing from the holoparasites Pae, Ocu, and Cau, including *CO, SVP, ELF4, AGL19* and *AGL16*, and *CDF1* and *CDF3* (Figure 6D). In terms of the regulatory pathways of flowering, Cau lost more genes related to the autonomous and hormone pathways than did Pae and Ocu, while Pae and Ocu lost more genes involved in the circadian and photoperiod pathways than did Cau. We suppose that this is due to the fact that Cau uses its host plant’s flowering signal FT to activate its own flowering (Shen et al., 2020), and that Pae and Ocu are unlikely to rely on light to signaling flowering, as they spend a significant amount of time living underground.

### The evolution of KAI2 gene family

After perceiving the strigolactones (SLs) released from the roots of adjacent host plants, the seeds of obligate parasites in the Orobanchaceae germinate (Cook et al., 1966; Yokota et al., 1998; Joel et al., 2013). In normal autotrophic plants, D14, an alpha/beta hydrolase protein, perceives SLs and regulates physiological processes related to development and stress resistance (Waters et al., 2017). In *Striga*, however, the receptor for SLs released by the host has been identified to be encoded by *KAI2d*. Importantly, *KAI2d* did not evolve from *D14*, but rather from the *KAI2* gene, which encodes the receptor of karrikin, through tandem duplication, large gene family expansion, and neofunctionalization (Conn et al., 2015; Yoshida et al., 2019).

We analyzed the *KAI2* gene families in the Orobanchaceae and in a few autotrophic species from lamiids, with *Arabidopsis* as the outgroup (Figure 7A). Only a single *KAI2* was identified in *Arabidopsis*, whereas in *Coffea canephora, Solanum lycopersicum*, and plants of the Orobanchaceae, at least two copies exist, suggesting that *KAI2* duplicated in the ancestor of lamiids resulting in the two paralogs, *KAI2c* and *KAI2i* (Figure 7A). We inspected the chromosomal fragments containing the *KAI2* gene in Llu and *Solanum lycopersicum*, and found that in both species, *KAI2c* and *KAI2i* are either on different chromosomes (Llu) or distantly located on the same chromosome (∼ 20 Mb in *Solanum lycopersicum*). Collinearity analysis indicated that the two chromosomal fragments containing the *KAI2* gene are obviously homologous with the fragment containing *KAI2* gene in *Arabidopsis* (Figure 7B). Given that no WGD events were detected in the ancestor of the lamiids ((Denoeud et al., 2014) and this study, Figure 2), the duplication of *KAI2* in the lamiids is a result of segmental duplication. *KAI2d* was then derived from duplication of *KAI2i* by tandem duplication (Figure 7A), as in Pja, Pcr, and Sas, *KAI2d* and *KAI2i* are adjacent (Figure 7B). In Pja and Pcr, the chromosome fragments containing *KAI2i* and *KAI2d* show good collinearity with the chromosome fragment containing *KAI2i* in Llu, while in Sas, Ocu, and Pae, the collinearity was not detected (Figure 7B). Thus, we inferred that this tandem duplication event, in which *KAI2d* came into existence, occurred in the common ancestor of the orobanchaceous parasitic plants. In Pja and Pcr the locations of these duplicated genes remained the same, whereas in Sas, Pae, and Ocu both genes translocated to other loci (Figure 7B).

**Figure 7.**
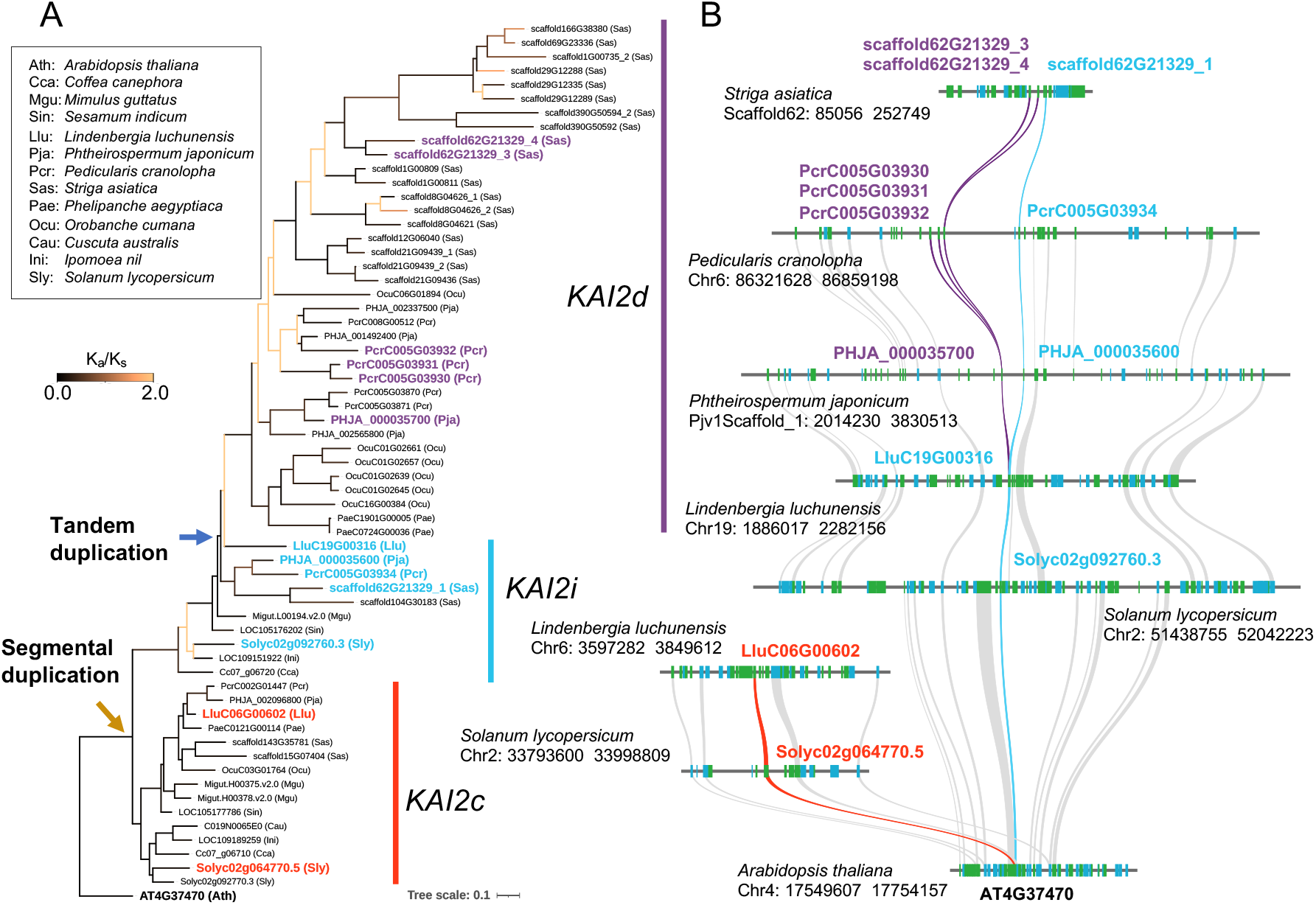
Molecular evolution of the *KAI2* gene family. (A) The phylogenetic tree of the *KAI2* family in 13 species. Blue and orange arrows indicate duplication events, and the colors of branches represent the values of K_a_/K_s_. (B) The collinearity relationships of chromosomal fragments containing the KAI2 gene in parasitic plants of the Orobanchaceae and closely related autotrophic species. Syntenic gene pairs are connected by curves. Curves connecting orthologs of *KAI2c, KAI2i*, and *KAI2d* genes are highlighted in different colors.

## Discussion

Orobanchaceae is the only family of plants containing species displaying all types of parasitic lifestyles, and it is preferred for studying the origin and evolution of parasitism. The phylogeny of the Orobanchaceae has been studied using plastome fragments (Young et al., 1999; Wolfe et al., 2005), ITS (Wolfe et al., 2005; McNeal et al., 2013), and individual nuclear genes (McNeal et al., 2013; Li et al., 2019). Even though most of these studies identified eight monophyletic clades, the phylogenetic relationships among these clades was not completely resolved. In this study, we assembled three genomes of the orobanchaceous plants *L. luchunensis, O. cumana*, and *Phelipanche aegyptiaca* (*P. aegyptiaca*). Together with the published/released transcriptome and genome data from other orobanchaceous plants, this study covered seven out of eight major clades of the Orobanchaceae. Using 907 nuclear OGs, we reconstructed a highly resolved and well-supported phylogeny of the Orobanchaceae, and the divergence times were estimated using a Bayesian molecular dating approach (Figure 1). Based on the phylogeny, we propose that 1) the common ancestor of the Orobanchaceae, an autotrophic plant, evolved parasitism (a haustorium) at least 38.58 Mya (95% HPD, 31.27-45.81 Mya); 2) the Cymbarieae, which comprises only facultative hemiparasites, was the earliest diverged parasitic lineage with low biodiversity; 3) by the Oligocene (33.90-23.03 Mya), a facultative hemiparasitic ancestor rapidly evolved into two lineages; one diverged into the Buchnereae (facultative hemiparasitic, obligate hemiparasitic, and holoparasitic species) and the Orobancheae (holoparasites), and the other evolved to be the Pedicularideae (facultative hemiparasitic species) and the Rhinantheae (facultative hemiparasitic and holoparasitic); these four clades are the most species-rich. Grasslands rapidly expanded during the Oligocene (Torsvik and Cocks, 2016), and this event may have provided orobanchaceous parasites with opportunities to exploit grasses and herbs, enabling the expansion of these parasites (Eriksson and Kainulainen, 2011; McNeal et al., 2013). Notably, contrasting with a recent study (Li et al., 2019), our scenario suggested that the hemiparasites evolved earlier than the holoparasites, which is consistent with the progressive evolution of increasing dependency of hosts.

Most species in the Orobanchaceae have 8 to 20 pairs of chromosomes, but the chromosome number can reach 132 in some species (Barker et al., 1988; Joel et al., 2013; Lyko and Wicke, 2021). Based on the variation in chromosome number, Lyko and Wicke (2021) proposed at least three ancient WGD events during the evolution of Orobanchaceae, including one WGD event in *Striga* and another in the Orobancheae clade. The WGD event in *Striga* was confirmed by a recent genome analysis (Yoshida et al., 2019). However, our collinearity analysis of the genome data from *L. luchunensis, Phtheirospermum japonicum* (*P. japonicum*), *Pedicularis cranolopha, Striga asiatica* (*S. asiatica*), *O. cumana*, and *P. aegyptiaca* did not reveal any other WGD events in the Orobanchaceae, except a single WGD event (*α*^*B*^) that occurred during the evolution of the ancestor of Buchnereae (32.64 ± 6.58 Mya) (Figure 2), the clade containing *S. asiatica*. Moreover, the 1:1 collinearity of the chromosomes of *L. luchunensis* and *O. cumana* (Supplemental Figure 5) also strongly suggests that no ploidization occurred after the divergence of *L. luchunensis* and *O. cumana*. When we inspected the emergence times of facultative parasitism, obligate parasitism, and later holoparasitism in the Orobanchaceae, no WGD events were found to have co-occurred with these evolutionary events (Figures 1 and 2), suggesting that genome duplications in the Orobanchaceae were not associated with the evolution of parasitism. However, by analyzing the evolutionary history of HHEGs in *P. japonicum, S. asiatica*, and *P. aegyptiaca*, we found that the *β*^*L*^ WGD event, which occurred long before the emergence of parasitism, was involved in the origin of at least 11% of the HHEGs. In all parasites, the retention rates of *β*^*L*^-duplicated genes are much higher among the HHEG than genome-wide (Figure 5B, Supplemental Table 10). Particularly, in the facultative hemiparasite *P. japonicum*, the retention rate of the *β*^*L*^ duplicated genes among the HHEGs was as high as 30.17%, far exceeding the expected retention rate under neutral selection (18.23%), indicating that these gene duplicates were retained by positive selection (Figure 5B). The gene duplicates caused by the *β*^*L*^ event may have allowed the ancestor of parasitic plants to acquire parasitism and a haustorium through neofunctionalization and/or subfunctionalization, and many of these genes remain being important in the lateral haustorium of facultative hemiparasitic plants. When the terminal haustorium/obligate parasitism evolved independently in the ancestors of *S. asiatica* and *P. aegyptiaca*, the HHEGs originated from the *β*^*L*^ seem to have become less important, indicated by the highly reduced retention rates of the *β*^*L*^-duplicated genes among the HHEGs. Notably, the ancestors of *S. asiatica* and *P. aegyptiaca* both experienced lineage-specific gene duplications in the HHEGs (Figure 5A), and these lineage-specific gene duplications are likely to be the driving force behind the evolution of the terminal haustorium and obligate parasitism.

The dodder *C. australis* and the endoparasite *Sapria himalayana* (*S. himalayana*) are both non-orobanchaceous species. *C. australis* has only 19671 genes (Sun et al., 2018), and it is among the higher plants with the fewest protein-coding genes. The *S. himalayana* genome comprises 55179 genes; however, most of them are transposon-like sequences, leaving only about 10000 low-copy genes (Cai et al., 2021). In contrast, the orobanchaceous parasites have many more genes than do *C. australis* or *S. himalayana* (Figure 4), and this is due to species-specific gene family expansions in different orobanchaceous parasites (Figure 4). Furthermore, of these expanded gene families, genes likely to be related to haustorium development and function were identified and many of the genes highly expressed in haustoria evolved from lineage-specific duplications (Figure 5). We propose that species-specifically expanded gene families may be the driving force underlying the adaptation of the parasites to different hosts.

Similar to *C. australis* and *S. himalayana*, large-scale gene loss was also detected in orobanchaceous parasites, especially in the holoparasites *O. cumana* and *P. aegyptiaca*. The reconstructed gene-loss history indicated that only 74 genes were missing in the common ancestor of the orobanchaceous parasites, when parasitism evolved; by contrast, 777 genes were deleted from the genome of the MRCA of *O. cumana* and *P. aegyptiaca*. Thus, gene loss became more severe as the degree of parasitism increased. Our gene loss data from the Orobanchaceae and the previous published data from *S. himalayana* (Cai et al., 2021) and the full mycoheterotrophic orchidaceous *Gastrodia elata* (Xu et al., 2021), which are all heterotrophs, all strongly support the idea that the degree of gene loss and parasitism are highly positively correlated. Accordingly, heterotrophic plants could be divided into four classes with increasing levels of heterotrophy and degrees of gene loss. In the first class are heterotrophs with 2-3% gene loss; these plants are facultative hemiparasites. In the second class are heterotrophs with 6-7% gene loss requiring hosts to complete certain developmental stages, and these are obligate hemiparasites and initial mycoheterotrophic plants, such as most species of Orchidaceae. In the third class are heterotrophs that have lost 13-15% genes, and these are holoparasites or full mycoheterotrophs with vegetative organs, such as *Cuscuta, Orobanche*, and *Gastrodia*. The fourth class comprises plants where gene loss is over 30%; these are endoparasites with only reproductive organs, such as *S. himalayana*. The lost genes in the heterotrophs of the first and second classes seem to be rather random, while the genes lost in the heterotrophs of the third class are related to photosynthesis, transcriptional regulation, stress adaptation, and organogenesis; the lost genes in the endoparasite *Sapria himalayana*, from the fourth class, includes almost all those lost in *P. aegyptiaca, O. cumana, C. australis*, and *Gastrodia elata*, representing an extreme form of parasitism.

Convergent gene loss in different parasitic plants may be the result of neutral evolution after the loss of purifying selection, as these genes are no longer required for parasitism. However, the loss of some genes may enhance the fitness of the parasitic plant. In *Striga*, the *protein phosphatase 2C* gene is mutated, leading to abscisic acid (ABA) insensitivity, with the impaired ABA signaling pathway leading to very high transpiration in *Striga*, and enabling this parasite to absorb nutrients from host xylem sap, even under drought conditions (Fujioka et al., 2019; Yoshida et al., 2019). The leafless *Cuscuta australis* has lost almost all the flowering regulatory pathways, and its *FT* gene, which encodes an important flowering signal, had pseudogenized; however, *C. australis* uses the FT proteins from host plants to activate its own flowering, enabling it to synchronize its flowering times with those of its hosts (Shen et al., 2020). In this manner, *C. australis* can parasitize a large number of host species with varying flowering times, and losing the flowering genes is thus an elegant adaptive strategy in *C. australis* (Shen et al., 2020). It would be interesting to study which pathways are affected by gene loss in orobanchaceous parasites and, importantly, the physiological and ecological impact incurred by gene loss.

Searcy and Maclnnis (1970) proposed a three-phase model of parasitic plant genome evolution: first, development of haustorium which enables a physical connection with host plants; secondly, after the haustorium has evolved, the parasites start to lose genes that are no longer needed, and thirdly, the parasites evolve to adapt to their specific hosts. The evolution of orobanchaceous parasites supports this model well. The *β*^*L*^ WGD event in the common ancestor of the Orobanchaceae and *Mimulus* generated many duplicated genes which likely contributed to the evolutionary gain of a haustorium; thereafter, gene loss events occurred in orobanchaceous parasites, especially in the holoparasitic species, and in parallel, expansions in specific gene families may have contributed to specialization and adaptation of the parasites to the hosts.

In this study, our phylogenomic analysis constructed a robust phylogenetic framework of the major lineages of the Orobanchaceae. Furthermore, genome evolution and comparative genomics analyses shed new light on the origin and evolution of parasitism in the Orobanchaceae. These genome resources and further molecular, genetic, and genomic studies on orobanchaceous species will provide many interesting insights into the evolution underlying the drastic physiological and morphological changes in these fascinating plants.

## Materials and Methods

### Estimation of genome size

Genome sizes were estimated based on flow cytometry and genomic *k*-mer analysis (Supplemental Table 1). A CyFlow Space-3000 (Partec, Germany) was used, and *Oryza sativa* L. ssp. *japonica* (370 Mb) and *Zea mays* ssp. *mays* var. B73 (2.1 Gb) were selected as the internal standards (Supplemental Table 1). In the genomic *k*-mer analysis, the genome sizes were estimated using ∼100 × coverage of short-read sequencing data (Supplemental Table 1), based on the *k*-mer (*k*=17) depth frequency distribution analysis embedded in the GCE software (v1.0.0) (Liu et al., 2020).

### Genome sequencing

Leaves of Llu and aboveground parts of stems of Ocu and Pae were harvested and cleaned with tap water before being used for DNA extraction. For short-read sequencing, DNA libraries were constructed using a VAHTS Universal Plus DNA Library Prep Kit for MGI (Vazyme, Nanjing, China) following the manufacturer’s instructions, and the libraries were sequenced on an MGI-SEQ 2000 (MGI Tech, Shenzhen, China) to generate 150-bp paired-end data for each species with approximately 100 × coverage. For long-read sequencing, we used the PromethION platform from Oxford Nanopore Technologies for Llu, and for Ocu and Pae, we used the PacBio Sequel II platform with the CLR mode. For each species, long-read data with approximately 100 × coverage were generated.

Tissues of Llu and Ocu were used for the Hi-C pipeline following a previously described method (Rao et al., 2014), including crosslinking, chromatin digestion with MboI (New England BioLabs, MA), labeling of DNA ends, ligation, purification, shearing, and biotin pull-down. The Hi-C libraries were sequenced on an MGI-SEQ 2000 (MGI Tech, Shenzhen, China) platform in the 150-bp paired-end mode, resulting in approximately 100 × coverage.

### Genome assembly and annotation

For genome assembly, we used the long-read data for *de novo* assembly, followed by error correction and polishing using the short-read data to obtain high-quality contigs. For Llu and Ocu, we used the Hi-C data to anchor the contigs onto chromosomes, to obtain chromosome-level assemblies (See Supplemental Methods for assembly software and procedures). To aid genome annotation, we obtained RNA-seq data from different tissues of orobanchaceous plants (Supplemental Table 3) using a DNBSEQ-T7 platform (MGI Tech, Shenzhen, China). Based on these data, we adopted an annotation pipeline integrating RNA-seq data, *ab initio* predicted features, and homologous sequence information to annotate repetitive sequences and protein-coding genes in the genomes (Supplemental Methods for details).

See Supplemental Methods for phylogenetic analysis and comparative genomics analysis.

## Data availability

The genome assembly, gene models, and sequence reads of Llu, Ocu, and Pae are available at the National Genomics Data Center (https://ngdc.cncb.ac.cn/) under BioProject number PRJCA008847. All data are also available from the corresponding author upon request.

## Acknowledgments

We thank Dr. Xiaopeng Yun (Inner Mongolia Academy of Agricultural and Animal Husbandry Sciences) for providing Ocu and Pae seeds, the Germplasm Bank of Wild Species in Southwest China for providing Llu seeds, and Dr. Steven Runo (Kenyatta University) for providing the photograph of *Striga*. Drs. Jun Zhao and Jian Zhang (Inner Mongolia Agricultural University) are thanked for the identification of Ocu. We are grateful to the High-Performance Computing Facility of the Department of Economic Plants and Biotechnology at the Kunming Institute of Botany for providing computing services. This study was supported by the National Natural Science Foundation of China (No. 32000179, Y.X.), the Strategic Priority Research Program of Chinese Academy of Sciences (No. XDPB16, J.W.), the CAS “Light of West China” Program (G.S.), the Special Research Assistant of Chinese Academy of Sciences (Y.X. and J.Z.), and the Postdoctoral Directional Training Foundation of Yunnan Province (Y.X.).

