## Supplemental Information for "Comparative genomics of orobanchaceous species with different parasitic lifestyles reveals the origin and stepwise evolution of plant parasitism"

**Supplemental Methods**

### Genome assembly

For Ocu and Pae, the subreads from the Pacbio sequencing were corrected, trimmed, and assembled by CANU (v2.2) (Koren et al., 2017). Contigs were polished with the PacBio data using GCpp (v1.9.0) (https://github.com/PacificBiosciences/gcpp) and then corrected with short-reads data using Pilon (v1.23) (Walker et al., 2014). For Llu, the long reads of ONT were de novo assembled using NextDenovo (v2.3.1) (https://github.com/Nextomics/NextDenovo), and the obtained contigs were corrected with NextPolish (v1.1.0) (Hu et al., 2020) using both short reads and long reads.

To obtain chromosome-scale assemblies for Llu and Ocu, we mapped the Hi-C data to contigs and constructed the Hi-C contact matrices, which were used to correct the assembly of contigs and build scaffolds using the 3D-DNA pipeline (v180922) (Dudchenko et al., 2017).

In order to evaluate the quality of the assembly, short reads were mapped to the assemblies using HISAT2 (Kim et al., 2019, 2), and the results from mapping were used for calling SNPs and indels using SAMtools (v1.9) (Li et al., 2009) and BCFtools (v1.9) (Danecek et al., 2021). The homozygous SNPs and indels were considered as assembly error, and thus the correct rates of assembly were calculated.

### Genome annotation

We first used the EDTA pipeline (v1.8.3) (Ou et al., 2019) to annotate repetitive sequences in the genomes, which integrates with the LTR_FINDER (Xu and Wang, 2007), LTRharvest (Ellinghaus et al., 2008), LTR_retriever (Ou and Jiang, 2018), Generic Repeat Finder (Shi and Liang, 2019), TIR-learner (Su et al., 2019), HelitronScanner (Xiong et al., 2014), and TEsorter (Zhang et al., 2019). EDTA first used these software to de novo annotate genome-wide transposons, and then the resulting transposons were integrated with the Dfam database (https://dfam.org/) into RepeatMasker (http://repeatmasker.org) to annotate repetitive elements.

To annotate the protein-coding genes, four steps were used to guide the annotation. 1) Augustus (v3.4.0) (Keller et al., 2011) for gene model prediction was trained with the MAKER-P pipeline (Campbell et al., 2014) using the RNA-seq data and the protein-coding genes in the close species (*Striga asiatica*, *Mimulus guttata*, *Arabidopsis thaliana*, *Solanum lycopersicum*, *Coffea canephora*, and *Cuscuta australis*). After three rounds of training of Augustus, de novo annotation of protein-coding genes was done. 2) The RNA-seq data were mapped to unmasked genomes using TopHat2 (v2.1.1) (Kim et al., 2013, 2), and the mapped reads were assembled to gene structures by Cufflinks (v2.2.1) (Trapnell et al., 2012); TRINITY (v2.8.5) (Haas et al., 2013) was used to de novo assemble the RNA-seq reads and thereafter PASA2 (v2.4.1) (Haas et al., 2003) was used to map the assembled contigs to the genomes and annotate the gene structures. These annotation results were merged. 3) The protein-coding genes in the closely related species were aligned to the unmasked genomes using GenBlastA (v1.0.1) (She et al., 2009), and GeneWise2 (v2.4.1) (Birney et al., 2004) was employed to generate gene structures. 4) The gene structures obtained from the above three steps were combined by EVidenceModeler (v1.1.1) (Haas et al., 2008). Then, the gene annotation results were compared with the repetitive element annotation results to remove the genes that overlap with repetitive elements. The final annotations were achieved.

**Phylogenetic analysis**

To resolve the phylogenetic relationships among the major clades of the Orobanchaceae, we collected data from 19 species of Orobanchaceae and three outgroup species (Supplementary Table 5). Six of the orobanchaceous species (*Lindenbergia luchunensis* (Llu), *Striga asiatica* (Sas), *Phtheirospermum japonicum* (Pja), *Pedicularis cranolopha* (Pcr), *Phelipanche aegyptiaca* (Pae), *Orobanche cumana* (Ocu)) and three outgroup species (*Coffea canephora*, *Mimulus guttatus* (Mgu), *Olea europaea*) have reference genomes, and for these species we directly used the amino acid sequences of all their gene models for the analysis. For the other 13 species of Orobanchaceae, we used their transcriptome data (see the Supplementary Table 5 for data sources). The transcriptome data were first *de novo* assembled by TRINITY (v2.8.5) (Haas et al., 2013), retaining the longest isoform of each unigene, searching for open reading frames and translating them into amino acid sequences using TransDecoder (v5.5.0) (https://github.com/TransDecoder/TransDecoder). The nine species with genomic data were first orthogrouped using OrthoFinder (v2.3.11) (Emms and Kelly, 2019) to obtain orthogroups (OGs). We followed the methodology of GeneFamilyClassifier pipeline in PlantTribes (Wall et al., 2008) to group the amino acid sequences of species with only transcriptome data into these OGs. Next, the OGs were screened following the criteria: 1) Sequences from genomic data are aligned and trimmed (using ClustalW2 (v2.1) (Larkin et al., 2007) default parameters for multiple sequence alignment, and thereafter trimAI (v1.2) (Capella-Gutiérrez et al., 2009) using gappyout mode for trimming), and the sequence lengths in the trimmed alignment matrix cannot be zero and gaps cannot exceed 50% of the whole matrix; 2) for the species whose genomes have been sequenced, each OG contains only a single gene from each species; 3) for the species whose genomes are not available, each OG contains maximum four sequences of a gene from each species; 4) for an individual OG, all species must have at least 1 sequence. In each eligible OG, the sequences identified from transcriptomes were added to the existing multiple sequence alignment matrix of that OG using the add function of MAFFT (v7.487) (Katoh and Standley, 2013), and in case of having multiple sequences from the same species, the consensus sequence were used for the matrix. The final number of OGs used for phylogenetic analysis was 907. For these OGs, we used both coalescent and concatenation methods for reconstruction of the phylogenetic tree. When the coalescent method was used, we first took the 907 OGs and built trees using RAxML-NG (v1.0.3) (model LG+G4, bootstrap 100) (Kozlov et al., 2019), and the resulted 907 trees were used for ASTRAL-III (v5.7.1) (Zhang et al., 2018) to construct the coalescent tree. When the concatenation method was used, we combined the multiple sequence alignment matrices of the 907 OGs to form a super matrix, which was further used for RAxML-NG (v1.0.3) (model LG+G4, bootstrap 100) (Kozlov et al., 2019) to construct the phylogenetic tree. The coalescent and concatenation methods yield an identical topology.

To estimate the divergence time for each node, we filtered the top 50 OGs with the smallest variance of the end node to root evolution distance. The CDS comparison matrix for these OGs was used for MCMCTREE in the PAML package (v4.9g) (Yang and Rannala, 2006), and due to lack of fossil evidence for Orobanchaceae, we used two secondary calibration points: the 95% highest posterior density limit of divergence time for Lamiales (70.56-99.84 Mya) and Orobanchaceae (37.56-63.46 Mya) (Xu et al., 2019).

**Identification of whole genome duplication (WGD) events**

WGD analysis was based on collinear fragments (CFs) between genomes. CFs are commonly obtained by analysis using blast-based identification of all homologs; however, CFs obtained in this manner include homologs generated by all the WGD events. Therefore, among the CFs identified by blast between two species, which recently evolved from a common ancestor, many CFs may be derived from certain WGD events before the speciation (Supplemental Figures 4A and B). If there were independent WGD events in any of these two species, the CFs resulted from the common WGD events before speciation will interfere with the identification of the recent species-specific WGD events happened after speciation, especially when the duplicated genes from ancient WGD are well retained, while those from the recent WGD events are rapidly lost (illustrated in Supplemental Figures 4A and B).

To identify the CFs generated from different WGD events, after obtaining all the homologs by orthogrouping, phylogenetic trees were constructed to detect the orthologs between the target and the reference species; the detected orthologs and their orders in the contigs were used for building CFs, which we named as the orthologous collinear fragments (OCFs) (Supplemental Figure 4C). OCFs are considered to be evolved after speciation, but not from the WGD events happened before their most recent common ancestor (MRCA). In this manner, for a given pair of species (A and B), the maximum depths of OCFs mapped to species B can be used to infer the WGD events in species A after A and B split from their MRCA, and vice versa (Supplemental Figure 4C).

Protein sequences of all gene models in every species were first clustered using OrthoFinder (v2.3.11) (Emms and Kelly, 2019), and thereafter multiple sequence alignment was performed for each obtained OGs using ClustalW2 (v2.1) (Larkin et al.,

2007), and gene trees were constructed using FastTree (v2.1.10) (Price et al., 2010). For a given pair of species, all orthologous genes were identified based on the topology of the gene tree. Using the orthologous gene pairs, collinear fragments were constructed with MCScanX_h (Wang et al., 2012), and these fragments were named OCFs. All species pairs among the selected species were analyzed using the above process to obtain all WGD events in the species tree.

**Estimation of timing of WGD events**

For each gene tree, all the binary nodes were traveled and determined whether the node is a "speciation node" or a "duplication node" based on if the two child nodes have common species (if yes, it is a "duplication node"; otherwise, it is a "speciation node"). Then we determined the phylogenetic placement of each node based on the MRCAs of the two child node's species.

To calculate the linear relationship between the divergence time and $K_s$, four speciation events (Figure 2, indicated by black circles) were chosen, as these events showed enough differences in divergence time. For each event, the "speciation nodes" corresponding to this event in all gene trees were filtered out. The $K_s$ values were calculated for all the child nodes under these "speciation nodes". Using SciPy (Virtanen et al., 2020), the $K_s$ values were used for determination of the smallest peak value based on Gaussian function (Supplementary Figure 5). Next, for the four speciation events (Figure 2, indicated by black circles) linear regression analysis was done using the peak values of $K_s$ and the divergence times estimated by molecular clock analysis (see Phylogenetic analysis), and the resulted parameters were used to infer the times of WGD events based on the peak values of $K_s$ obtained from the "duplication nodes".

**Reconstruction of ancestral karyotypes**

To reconstruct the ancestral karyotypes of Orobanchaceae, we used the Llu, Pcr, and Ocu genomes, which have chromosome-level assembly, and the genome of the outgroup Mgu. In identification the WGD events, we obtained all the OCFs of Mgu with Llu, Pcr, and Ocu, respectively. The anchored genes on the OCFs were used for reconstructing the ancestral karyotypes. The order of these orthologous genes on the chromosomes of these four genomes and the species trees of the four species were fed to the MGRA software (v2.3) (Alekseyev and Pevzner, 2009) to generate the ancestral karyotypes of each node in the species tree.

**Expansion and contraction of gene families**

To determine whether significant expansion and contraction of gene families at each evolutionary node in the species tree, we used a pipeline named EasyBadiRate (https://github.com/SouthernCD/EasyBadiRate). In this pipeline, for each gene family, we used the free model in BadiRate (v1.35) (Librado et al., 2012) to estimate the sizes of the ancestral gene families in all clades of the species tree. For the branches whose gene families did not experience family size changes, they were set to be the background branches, which have the same family turnover rate; thereafter, based on this model, we re-estimated the likelihood, and the model was regarded as the null hypothesis. For branches that experienced size changes, an alternative hypothesis for each branch was built by forcing the given branch to follow the same turnover rate with the background branches. A branch that experienced size changes was significant, if AIC (alternative hypothesis) - AIC (null hypothesis) > 2 (Akaike's information criterion (AIC) was computed from the likelihood and numbers of parameters in each model) (Burnham and Anderson, 2004). Hereafter, the significantly expanded and contracted gene families in all lineages were obtained.

To compare the sizes of gene families across species, we followed a previously published statistic $F$-index (Sun et al., 2018; Xu et al., 2021), which describe the size differences of conserved gene families in all autotrophs.

$$F_{ij} = \frac{\log_2\left(a\frac{c_{ij}}{N_j} + \frac{1}{2}\right) + 1}{\log_2\left(a + \frac{1}{2}\right) + 1}$$

$$a = \frac{S^2 - 2S}{2}$$

In this formula, $i$ refers to a given species and $j$ refers to a given family, $c_{ij}$ is the number of genes in $j$ family in $i$ species, and $N_j$ is the total number of genes in $j$ family of all species, and $S$ represents the number of species. $F$-index is a modified log2-fold-change coefficient ($\log_2 \frac{c_{ij}}{N_j}$), so the relationship between $F$-index and $\log_2 \frac{c_{ij}}{N_j}$ is linear. $F$-indices range from 0 to 1, and when $F_{ij} = 0.5$, in species $i$, the gene number in gene family $j$ equals to the average size of this gene family in all species; if $F_{ij} = 0$, there are no genes in family $j$ in species $i$, and if $F_{ij} = 1$, it indicates that only species $i$, but not the others, harbors the gene family $j$.

**Identification of gene loss**

To investigate the gene loss events occurred in the genomes of orobanchaceous parasitic plants and *Cuscuta australis* (Cau), we used the protein-coding genes of these species and a few closely related autotrophs for phylogenomic analysis. The species used in the analysis were five orobanchaceous parasitic plants (Pae, Ocu, Sas, Pja, Pcr), one orobanchaceous autotrophic plant (Llu), two species of Lamiales (Mgu, *Sesamum indicum*), parasitic plant Cau from Convolvulaceae, one autotrophic plant from Convolvulaceae (*Ipomoea nil* (Ini)), two species from other order in lamiids (*Solanum lycopersicum* and *Coffea canephora*), and the outgroup species *Arabidopsis thaliana* (Ath).

First, we clustered the amino acid sequences of gene models from these 13 species by OrthoFinder (v2.3.11) (Emms and Kelly, 2019), and after constructing the gene trees by TreeBeST (v1.9.2) (Vilella et al., 2009), we obtained the orthogroups (OGs) from the gene trees. The conserved OGs were selected (a "conserved" OG must have gene(s) from the Ath, Llu (closely autotrophs for orobanchaceous parasitic plants) and Ini (closely autotrophs for Cau), and at least 3 other 4 autotrophs species). We finally obtained a total of 11,337 conserved OGs. By counting the missing genes in the parasitic plants, the candidate OGs with gene loss for each species were obtained.

False positives in the identification of gene loss can occur, due to errors of gene annotation, orthogrouping, and phylogenetic analysis. To minimize this possibility, we adopted a pipeline by Sun et al. (Sun et al., 2018) and Xu et al. (Xu et al., 2021) with modifications:

1. If the orthologous gene of a parasitic plant is missing in an OG, we assigned the homolog (amino acid sequence) from the autotroph to the genome of the parasitic plant using GenBlastA (v1.0.1) (She et al., 2009). If the aligned region does not contain a known gene (less than 80% overlap), the region is re-annotated using GeneWise2 (v2.4.1) (Birney et al., 2004) to obtain possible missing genes from annotation.

2. For each conserved OG, multiple sequence alignment was performed using the gene sequences from the autotrophs in that OG, and the profile hidden Markov models (HMM) of that OG was constructed using HMMER (v3.1b2) (Eddy, 2011). The complete set of annotated genes of the parasitic plant and the missing genes obtained in the previous step were combined into one dataset, and each sequence was compared to the HMM profile of each OG using HMMER (v3.1b2) (Eddy, 2011) to find the best OG for each sequence (E-value < 1e-20, coverage > 50%).

3. If an OG still lack of the ortholog from parasitic plants, the lost ortholog is considered to be genuine.

After identifying gene loss events in all parasitic plants (Dataset S3), we considered that the genes commonly lost among multiple species were lost in their MRCA. Based on this scenario, we assigned the numbers of gene loss events to the branches of species tree (Figure 6A).

**Identification of haustorially highly expressed genes**

We used the transcriptome data (Supplemental Table 7) from Pja, Sas, and Pae to identify haustorially highly expressed genes in these species. After mapping the transcriptome data to the respective reference genome by HISAT2 (Kim et al., 2019, 2), we obtained the counts of each gene using featureCounts (Liao et al., 2014), which were next normalized using edgeR (Robinson et al., 2010), and the differentially expressed genes (DEGs) between different samples were obtained ($|\log2(\text{fold change})| \geq 1$ and p-value $\leq$ 0.01). For each species, transcriptomes generated from haustorial tissues were grouped into a haustorium group and the rest were grouped into a non-haustorium group. After removing redundancy, DEGs whose relative expression levels are greater in the haustorium group than in the non-haustorium group were defined as the haustorially highly expressed genes.

**Exponential model of gene loss after WGD events**

According to the Ren et al. (2018), in angiosperms, the number of gene duplicates retained in the genome after a WGD event decreases exponentially.

$$R = a^k + b$$

Where $R$ is the percentage of retained gene duplicates, $k$ is $K_s$ peak value of the duplicated genes from WGD event, and $a$ and $b$ are constants. Comparison of the 22 known WGDs in angiosperms revealed that the half-life of gene duplicates in the form of $K_s$ is approximately 0.34 (Ren et al., 2018). Accordingly, when $R$ = 1 and 0.5, $k$ = 0 and 0.34, respectively. It can be solved that $a$ = 0.13020 and $b$ = 0. The formula is:

$$R = 0.13020^k$$

**Nucleic acid extraction**

Genomic DNA was extracted from Llu, Ocu and Pae using QIAamp DNA Mini Kit/DNeasy Plant Mini Kit. The integrity of the DNA was determined with the Agilent 4200 Bioanalyzer (Agilent Technologies, Palo Alto, California). Eight micrograms of genomic DNA were sheared using g-Tubes (Covaris), and concentrated with AMPure PB magnetic beads. To aid genome annotation, we obtained RNA-seq data from different tissues of orobanchaceous plants (Supplemental Table 3). The total RNA was extracted from these tissues using the TRI Reagent (Sigma). The MGIEasy RNA library preparation kit was used for library construction.

**Supplemental information References**


**Alekseyev, M. A., and Pevzner, P. A.** (2009). Breakpoint graphs and ancestral genome reconstructions. *Genome Res.* **19**:943–957.

**Becher, H., Brown, M. R., Powell, G., Metherell, C., Riddiford, N. J., and Twyford, A. D.** (2020). Maintenance of Species Differences in Closely Related Tetraploid Parasitic Euphrasia (Orobanchaceae) on an Isolated Island. *Plant Communications* **1**:100105.

**Birney, E., Clamp, M., and Durbin, R.** (2004). GeneWise and Genomewise. *Genome Res* **14**:988–995.

**Boachon, B., Buell, C. R., Crisovan, E., Dudareva, N., Garcia, N., Godden, G., Henry, L., Kamileen, M. O., Kates, H. R., Kilgore, M. B., et al.** (2018). Phylogenomic Mining of the Mints Reveals Multiple Mechanisms Contributing to the Evolution of Chemical Diversity in Lamiaceae. *Molecular Plant* **11**:1084–1096.

**Burnham, K. P., and Anderson, D. R. eds.** (2004). *Model Selection and Multimodel Inference*. New York, NY: Springer New York.

**Capella-Gutiérrez, S., Silla-Martínez, J. M., and Gabaldón, T.** (2009). trimAl: a tool for automated alignment trimming in large-scale phylogenetic analyses. *Bioinformatics* **25**:1972–1973.

**Chen, J., Yu, R., Dai, J., Liu, Y., and Zhou, R.** (2020). The loss of photosynthesis pathway and genomic locations of the lost plastid genes in a holoparasitic plant Aeginetia indica. *BMC Plant Biology* **20**:199.

**Chen, L., Guo, Q., Zhu, Z., Wan, H., Qin, Y., and Zhang, H.** (2021). Integrated analyses of the transcriptome and small RNA of the hemiparasitic plant Monochasma savatieri before and after establishment of parasite-host association. *BMC Plant Biol* **21**:90.

**Cui, S., Kubota, T., Nishiyama, T., Ishida, J. K., Shigenobu, S., Shibata, T. F., Toyoda, A., Hasebe, M., Shirasu, K., and Yoshida, S.** (2020). Ethylene signaling mediates host invasion by parasitic plants. *Science Advances* Advance Access published October 2020, doi:10.1126/sciadv.abc2385.

**Denoeud, F., Carretero-Paulet, L., Dereeper, A., Droc, G., Guyot, R., Pietrella, M., Zheng, C., Alberti, A., Anthony, F., Aprea, G., et al.** (2014). The coffee genome provides insight into the convergent evolution of caffeine biosynthesis. *Science* **345**:1181–1184.

**Eddy, S. R.** (2011). Accelerated Profile HMM Searches. *PLOS Computational Biology* **7**:e1002195.

**Emms, D. M., and Kelly, S.** (2019). OrthoFinder: phylogenetic orthology inference for comparative genomics. *Genome Biol* **20**:238.

**Haas, B. J., Papanicolaou, A., Yassour, M., Grabherr, M., Blood, P. D., Bowden, J., Couger, M. B., Eccles, D., Li, B., Lieber, M., et al.** (2013). De novo transcript sequence reconstruction from RNA-seq using the Trinity platform for reference generation and analysis. *Nat Protoc* **8**:1494–1512.

**Hellsten, U., Wright, K. M., Jenkins, J., Shu, S., Yuan, Y., Wessler, S. R., Schmutz, J., Willis, J. H., and Rokhsar, D. S.** (2013). Fine-scale variation in meiotic recombination in *Mimulus* inferred from population shotgun sequencing. *Proc. Natl. Acad. Sci. U.S.A.* **110**:19478–19482.

**Kado, T., and Innan, H.** (2018). Horizontal Gene Transfer in Five Parasite Plant Species in Orobanchaceae. *Genome Biology and Evolution* **10**:3196–3210.

**Katoh, K., and Standley, D. M.** (2013). MAFFT Multiple Sequence Alignment Software Version 7: Improvements in Performance and Usability. *Molecular Biology and Evolution* **30**:772–780.

**Kim, D., Paggi, J. M., Park, C., Bennett, C., and Salzberg, S. L.** (2019). Graph-based genome alignment and genotyping with HISAT2 and HISAT-genotype. *Nat Biotechnol* **37**:907–915.

**Kozlov, A. M., Darriba, D., Flouri, T., Morel, B., and Stamatakis, A.** (2019). RAxML-NG: a fast, scalable and user-friendly tool for maximum likelihood phylogenetic inference. *Bioinformatics* **35**:4453–4455.

**Larkin, M. A., Blackshields, G., Brown, N. P., Chenna, R., McGettigan, P. A., McWilliam, H., Valentin, F., Wallace, I. M., Wilm, A., Lopez, R., et al.** (2007). Clustal W and Clustal X version 2.0. *Bioinformatics* **23**:2947–2948.

**Liao, Y., Smyth, G. K., and Shi, W.** (2014). featureCounts: an efficient general purpose program for assigning sequence reads to genomic features. *Bioinformatics* **30**:923–930.

**Librado, P., Vieira, F. G., and Rozas, J.** (2012). BadiRate: estimating family turnover rates by likelihood-based methods. *Bioinformatics* **28**:279–281.

**Ma, L., Dong, C., Song, C., Wang, X., Zheng, X., Niu, Y., Chen, S., and Feng, W.** (2021). De novo genome assembly of the potent medicinal plant Rehmannia glutinosa using nanopore technology. *Computational and Structural Biotechnology Journal* **19**:3954–3963.

**Marx, H. E., Carboni, M., Douzet, R., Perrier, C., Delbart, F., Thuiller, W., Lavergne, S., and Tank, D. C.** (2021). Can functional genomic diversity provide novel insights into mechanisms of community assembly? A pilot study from an invaded alpine streambed. *Ecology and Evolution* **11**:12075–12091.

**Price, M. N., Dehal, P. S., and Arkin, A. P.** (2010). FastTree 2 – Approximately Maximum-Likelihood Trees for Large Alignments. *PLOS ONE* **5**:e9490.

**Rao, G., Zhang, J., Liu, X., Lin, C., Xin, H., Xue, L., and Wang, C.** (2021). De novo assembly of a new Olea europaea genome accession using nanopore sequencing. *Hortic Res* **8**:64.

**Robinson, M. D., McCarthy, D. J., and Smyth, G. K.** (2010). edgeR: a Bioconductor package for differential expression analysis of digital gene expression data. *Bioinformatics* **26**:139.

**She, R., Chu, J. S.-C., Wang, K., Pei, J., and Chen, N.** (2009). GenBlastA: enabling BLAST to identify homologous gene sequences. *Genome Res* **19**:143–149.

**Sun, G., Xu, Y., Liu, H., Sun, T., Zhang, J., Hettenhausen, C., Shen, G., Qi, J., Qin, Y., Li, J., et al.** (2018). Large-scale gene losses underlie the genome evolution of parasitic plant Cuscuta australis. *Nat Commun* **9**:2683.

**Vilella, A. J., Severin, J., Ureta-Vidal, A., Heng, L., Durbin, R., and Birney, E.** (2009). EnsemblCompara GeneTrees: Complete, duplication-aware phylogenetic trees in vertebrates. *Genome Res* **19**:327–335.

**Virtanen, P., Gommers, R., Oliphant, T. E., Haberland, M., Reddy, T., Cournapeau, D., Burovski, E., Peterson, P., Weckesser, W., Bright, J., et al.** (2020). SciPy

1.0: fundamental algorithms for scientific computing in Python. *Nat Methods* **17**:261–272.

**Wall, P. K., Leebens-Mack, J., Müller, K. F., Field, D., Altman, N. S., and dePamphilis, C. W.** (2008). PlantTribes: a gene and gene family resource for comparative genomics in plants. *Nucleic Acids Res* **36**:D970–D976.

**Wang, Y., Tang, H., DeBarry, J. D., Tan, X., Li, J., Wang, X., Lee, T., Jin, H., Marler, B., Guo, H., et al.** (2012). MCScanX: a toolkit for detection and evolutionary analysis of gene synteny and collinearity. *Nucleic Acids Res* **40**:e49.

**Wertheim, J. O., Murrell, B., Smith, M. D., Kosakovsky Pond, S. L., and Scheffler, K.** (2015). RELAX: Detecting Relaxed Selection in a Phylogenetic Framework. *Molecular Biology and Evolution* **32**:820–832.

**Westwood, J. H., dePamphilis, C. W., Das, M., Fernández-Aparicio, M., Honaas, L. A., Timko, M. P., Wafula, E. K., Wickett, N. J., and Yoder, J. I.** (2012). The Parasitic Plant Genome Project: New Tools for Understanding the Biology of *Orobanche* and *Striga*. *Weed sci.* **60**:295–306.

**Wickett, N. J., Honaas, L. A., Wafula, E. K., Das, M., Huang, K., Wu, B., Landherr, L., Timko, M. P., Yoder, J., Westwood, J. H., et al.** (2011). Transcriptomes of the parasitic plant family Orobanchaceae reveal surprising conservation of chlorophyll synthesis. *Curr Biol* **21**:2098–2104.

**Xu, W.-Q., Losh, J., Chen, C., Li, P., Wang, R.-H., Zhao, Y.-P., Qiu, Y.-X., and Fu, C.-X.** (2019). Comparative genomics of figworts ( *Scrophularia* , Scrophulariaceae), with implications for the evolution of *Scrophularia* and Lamiales: Comparative genomics of *Scrophularia*. *Jnl of Sytematics Evolution* **57**:55–65.

**Xu, Y., Lei, Y., Su, Z., Zhao, M., Zhang, J., Shen, G., Wang, L., Li, J., Qi, J., and Wu, J.** (2021). A chromosome-scale *Gastrodia elata* genome and large-scale comparative genomic analysis indicate convergent evolution by gene loss in mycoheterotrophic and parasitic plants. *The Plant Journal* **108**:1609–1623.

**Yang, Z., and Rannala, B.** (2006). Bayesian estimation of species divergence times under a molecular clock using multiple fossil calibrations with soft bounds. *Mol Biol Evol* **23**:212–226.

**Yang, Z., Wafula, E. K., Honaas, L. A., Zhang, H., Das, M., Fernandez-Aparicio, M., Huang, K., Bandaranayake, P. C. G., Wu, B., Der, J. P., et al.** (2015). Comparative Transcriptome Analyses Reveal Core Parasitism Genes and Suggest Gene Duplication and Repurposing as Sources of Structural Novelty. *Molecular Biology and Evolution* **32**:767–790.

**Yoshida, S., Kim, S., Wafula, E. K., Tanskanen, J., Kim, Y.-M., Honaas, L., Yang, Z., Spallek, T., Conn, C. E., Ichihashi, Y., et al.** (2019). Genome sequence of Striga asiatica provides insight into the evolution of plant parasitism. *Curr Biol* **29**:3041-3052.e4.

**Zhang, C., Rabiee, M., Sayyari, E., and Mirarab, S.** (2018). ASTRAL-III: polynomial time species tree reconstruction from partially resolved gene trees. *BMC Bioinformatics* **19**:153.

**Zhang, X., Li, C., Wang, L., Fei, Y., and Qin, W.** (2019). Analysis of Centranthera grandiflora Benth Transcriptome Explores Genes of Catalpol, Acteoside and Azafrin Biosynthesis. *International Journal of Molecular Sciences* **20**:6034.

**Zhang, C., Zhang, T., Luebert, F., Xiang, Y., Huang, C.-H., Hu, Y., Rees, M., Frohlich, M. W., Qi, J., Weigend, M., et al.** (2020). Asterid Phylogenomics/Phylotranscriptomics Uncover Morphological Evolutionary Histories and Support Phylogenetic Placement for Numerous Whole-Genome Duplications. *Molecular Biology and Evolution* **37**:3188–3210.

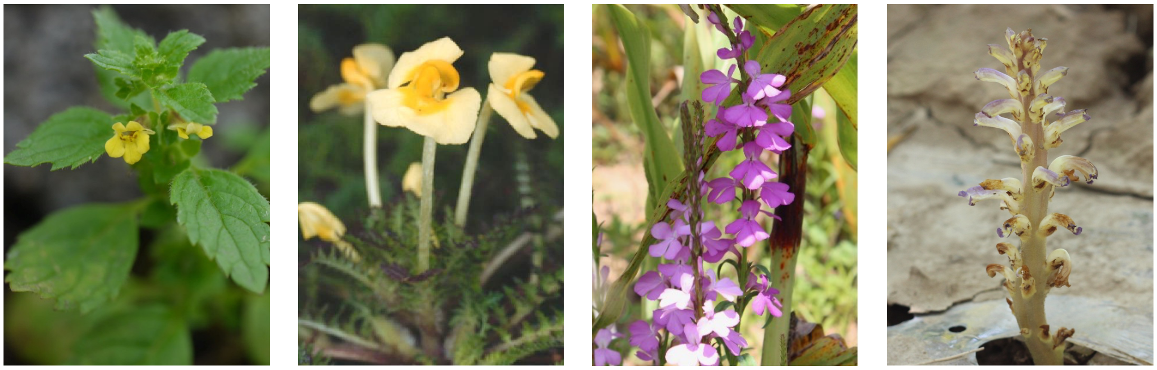

***Lindenbergia*** Autotrophic

***Pedicularis*** Facultative Hemiparasitic

***Striga*** Obligate Hemiparasitic

***Orobanche*** Obligate Holoparasitic

**Figure S1.** Plant species in the family Orobanchaceae spanning from non-parasitic to increasing levels of parasitism.

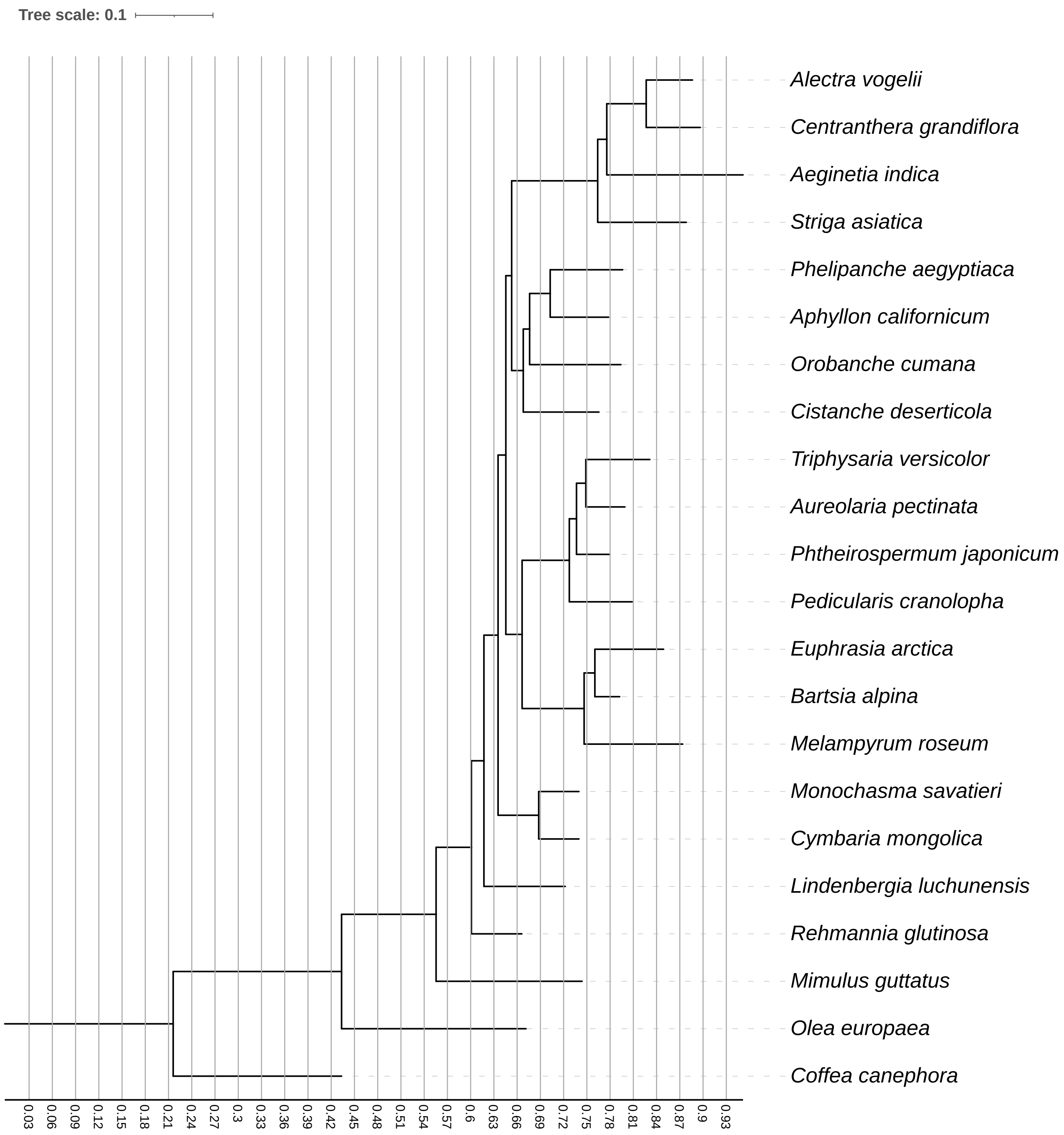


**Figure S2.** Phylogenetic tree was constructed using low-copy orthogroups from 19 orobanchaceous species and three outgroup plants. (Coalescent and concatenation method yielded identical topologies with 100% support for all the nodes, branch lengths were calculated by concatenation method. See Phylogenetic analysis)

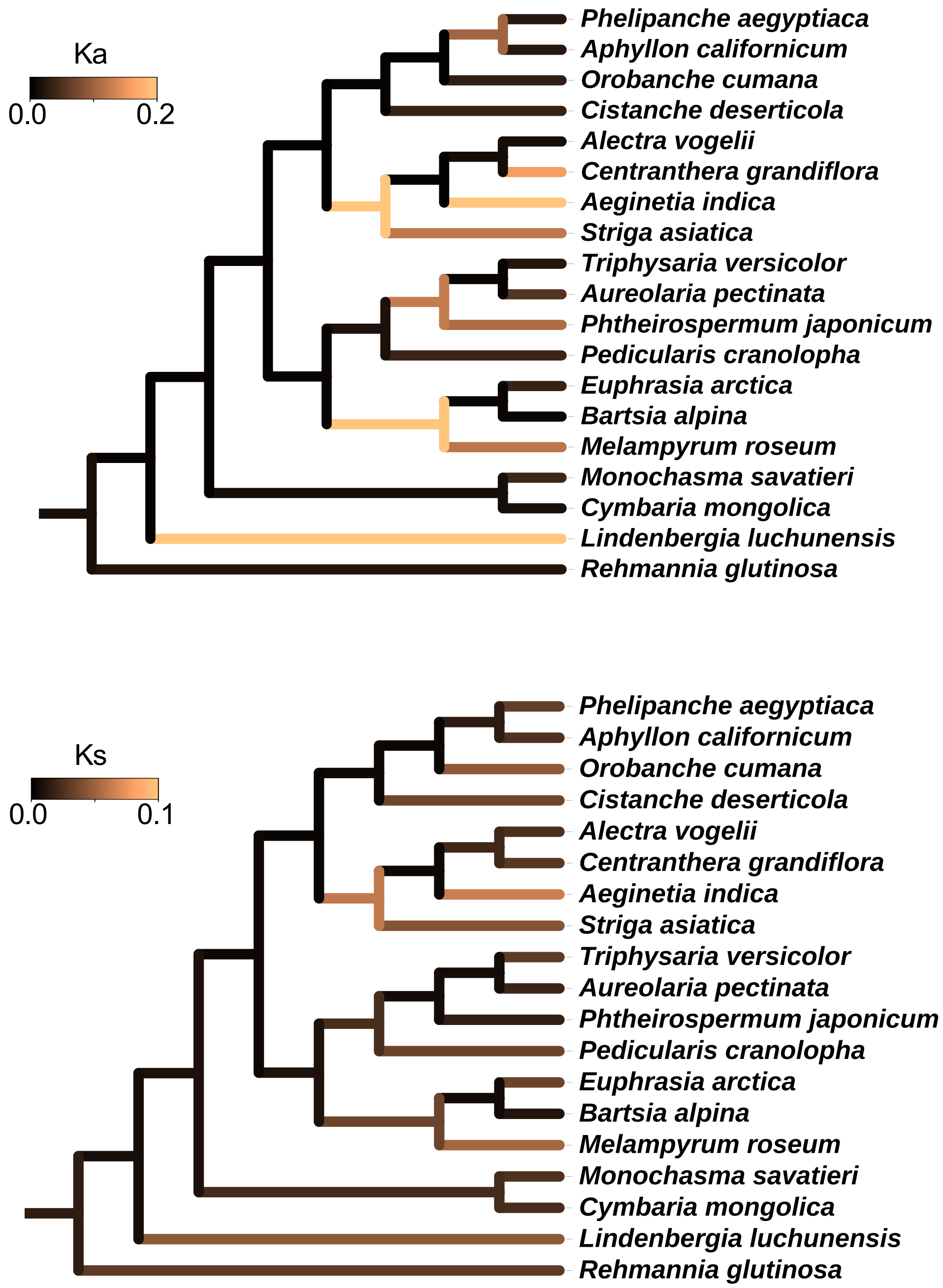


**Figure S3.** Evolution of $K_a$ and $K_s$ shown on the orobanchaceous phylogeny.

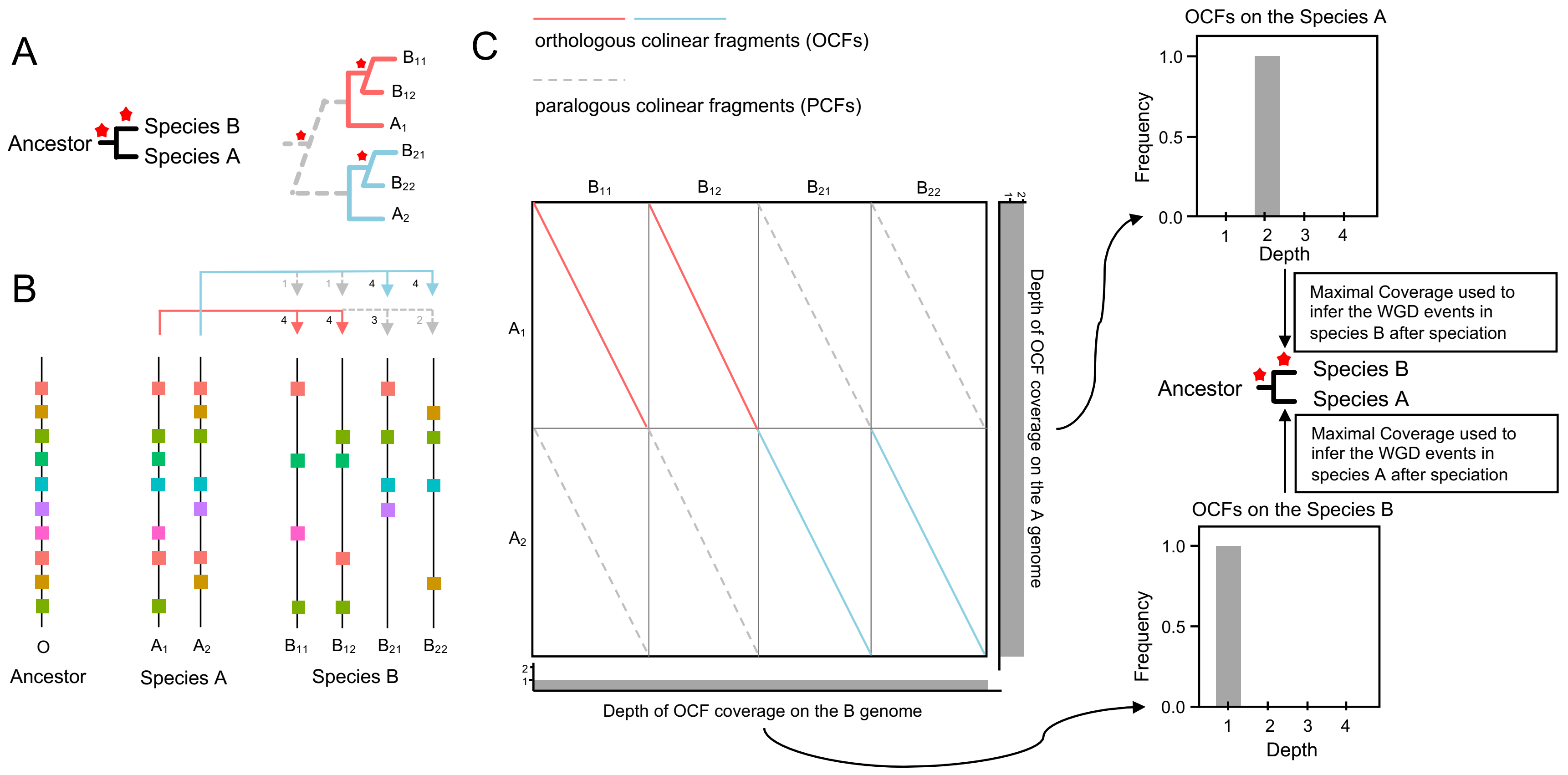


**Figure S4.** Identification of WGD events based on OCFs.

**(A)** Schematic illustration of a case in which two WGD events happened relatively closely. Species A and species B share a common WGD event before their MRCA, and species B independently experienced a WGD event after speciation. The figure on the right illustrate the evolutionary history of the homologous chromosomes under this scenario, assuming the “ancestor” had only one chromosome.

**(B)** Illustration of gene loss in the species A and B under the evolutionary scenario shown in (A). Although WGD events are usually followed by strong gene loss, colinear fragments can still be found on homologous chromosomes between species A and B. The anchor gene pairs are more easily found on homologous chromosomes caused by the younger WGD (red and blue arrows), while the anchor gene pairs are more difficult to identify from the older WGD (gray arrows). Colored blocks represent genes, and numbers next to the arrows represent the numbers of anchor gene pairs between homologous chromosomes.

**(C)** CFs between species A and B. Based on the phylogenetic relationships of anchor genes, CFs (colinear fragments) can be distinguished into orthologous colinear fragments (OCFs) and paralogous colinear fragments (PCFs). Counting the depth of OCF coverage on the A and B genomes can be used to determine the WGD events that occurred independently in species A and B after speciation from their MRCA. The depths of OCF coverage on the A and B genome can be displayed as a pair of histogram plots, as shown in Figure 2.

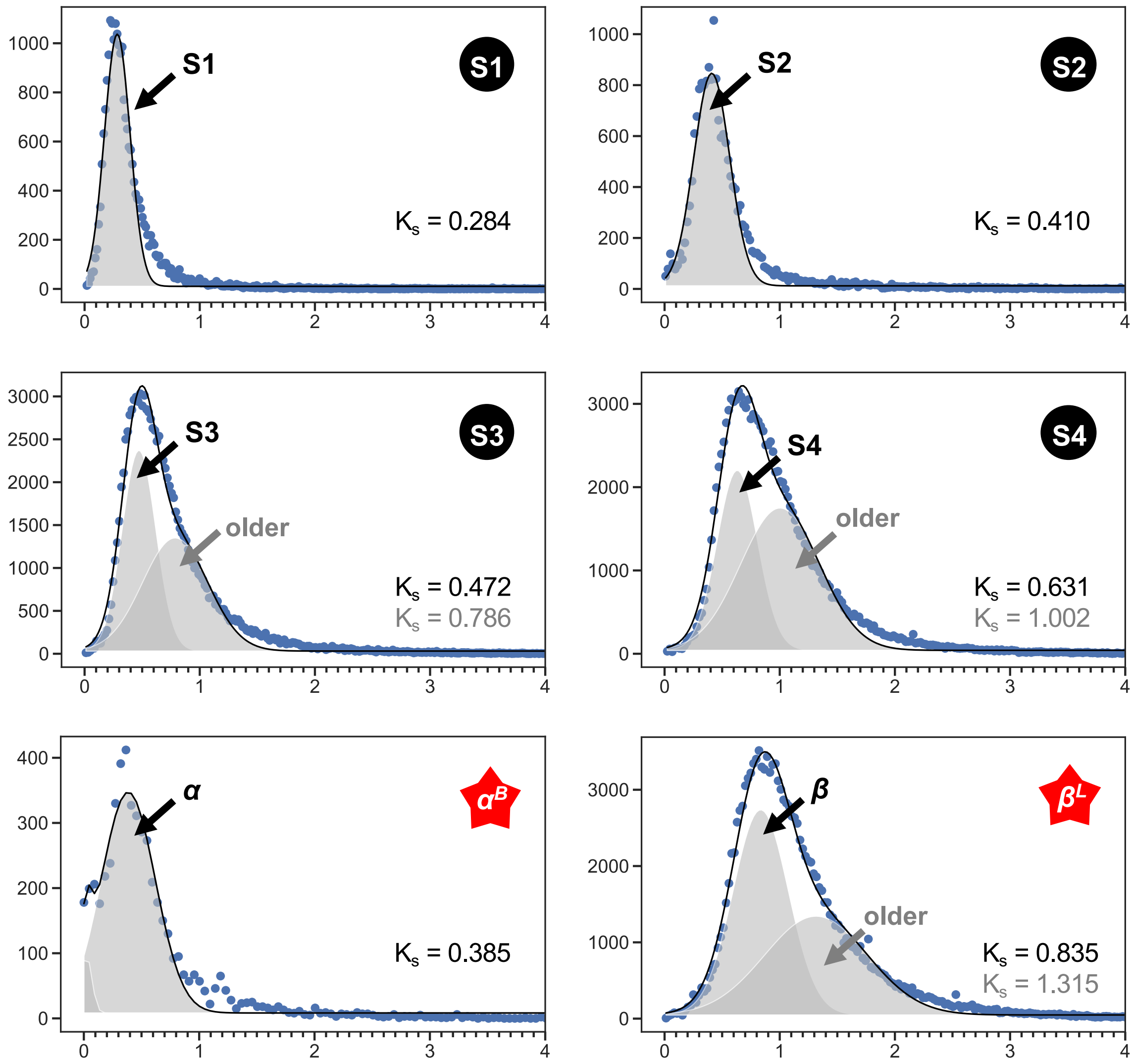


**Figure S5.** $K_s$ distribution and Gaussian fit of speciation events and WGD events. The blue dots are observed data, the black line is the fitted curve, and the gray shading is the Gaussian distribution.

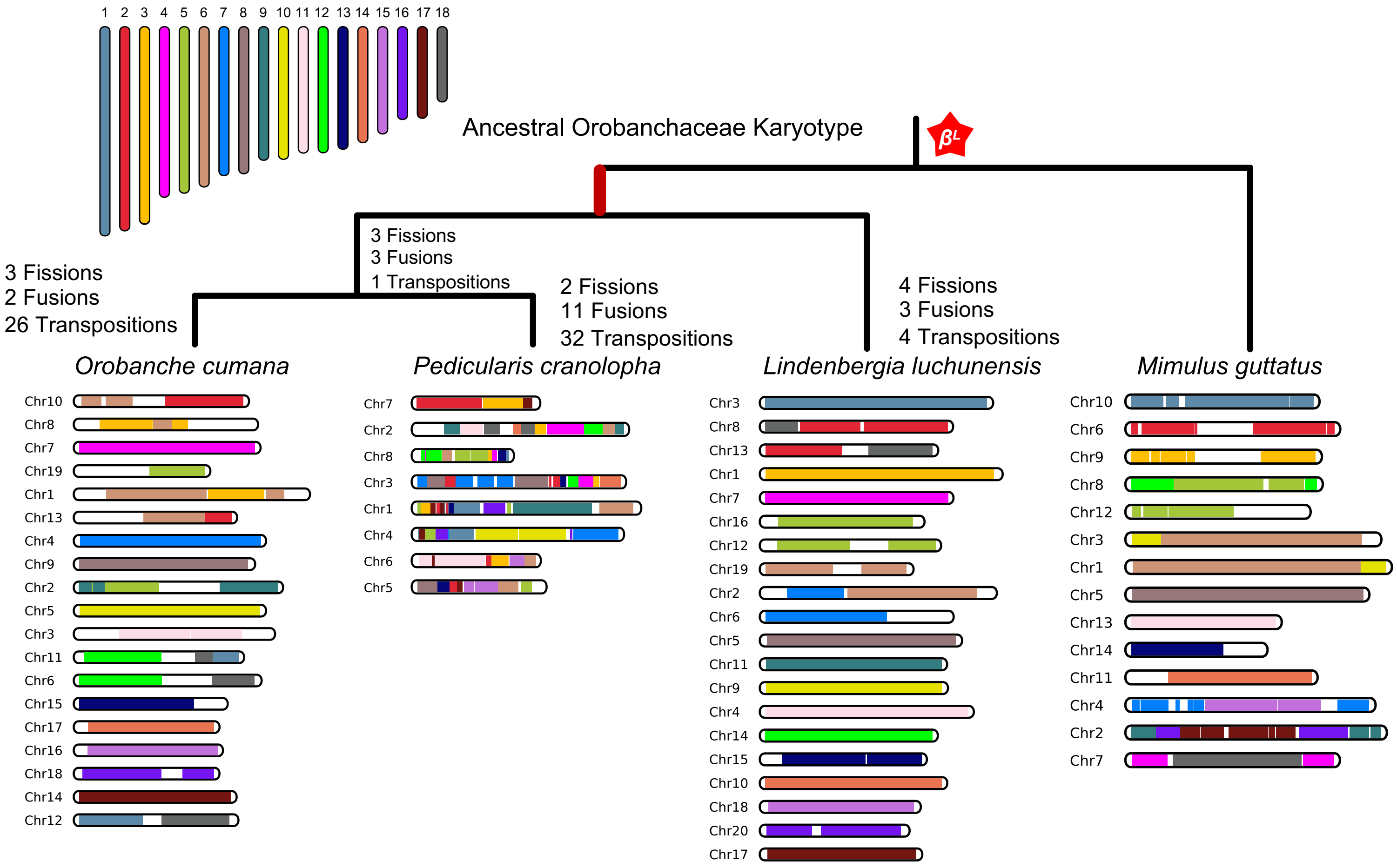


**Figure S6.** Eighteen ancestral Orobanchaceae protochromosomes were constructed using *Mimulus guttatus* as an outgroup.

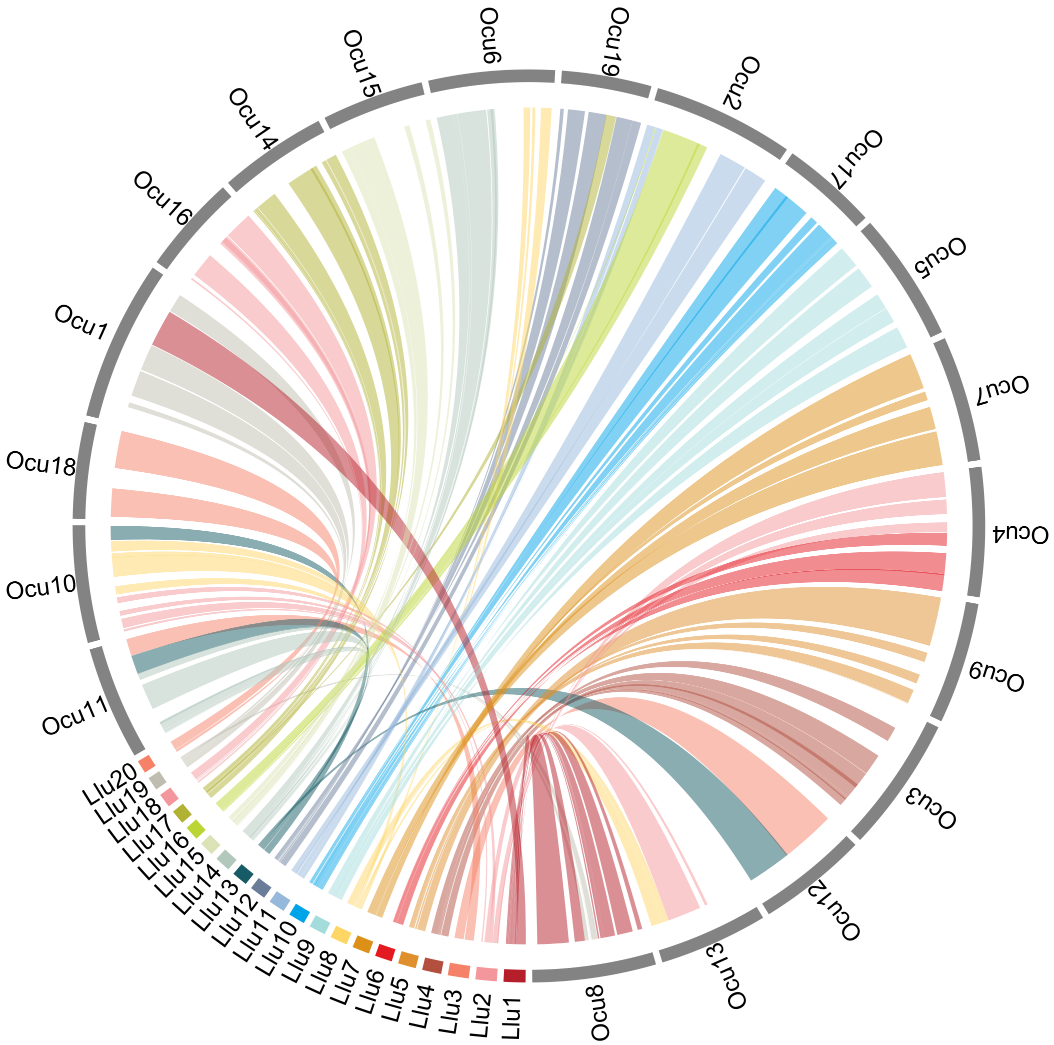


**Figure S7.** Circos plots showing conserved synteny between *Lindenbergia luchunensis* (Llu) and *O. cumana* (Ocu).

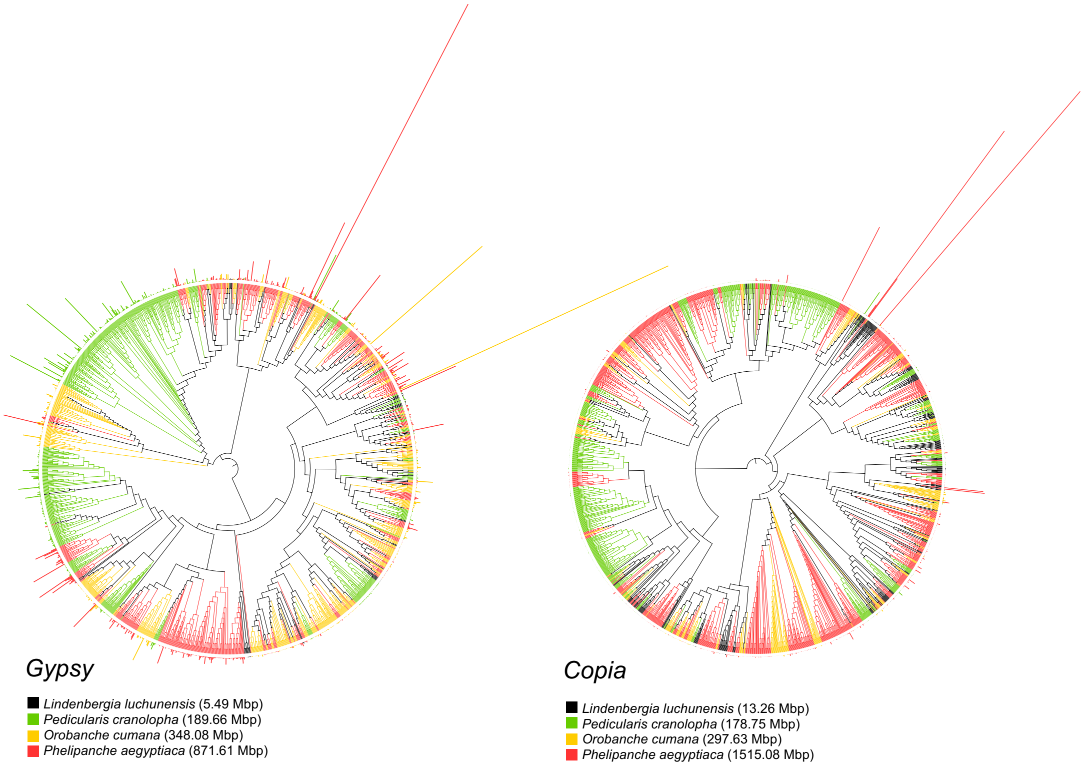


**Figure S8.** Phylogeny of LTR/Gypsy and LTR/Copia in *Lindenbergia luchunensis*, *Pedicularis cranolopha*, *Orobanche cumana*, and *Phelipanche aegyptiaca*. Bars outside dendrogram denote copy numbers of the repetitive family.

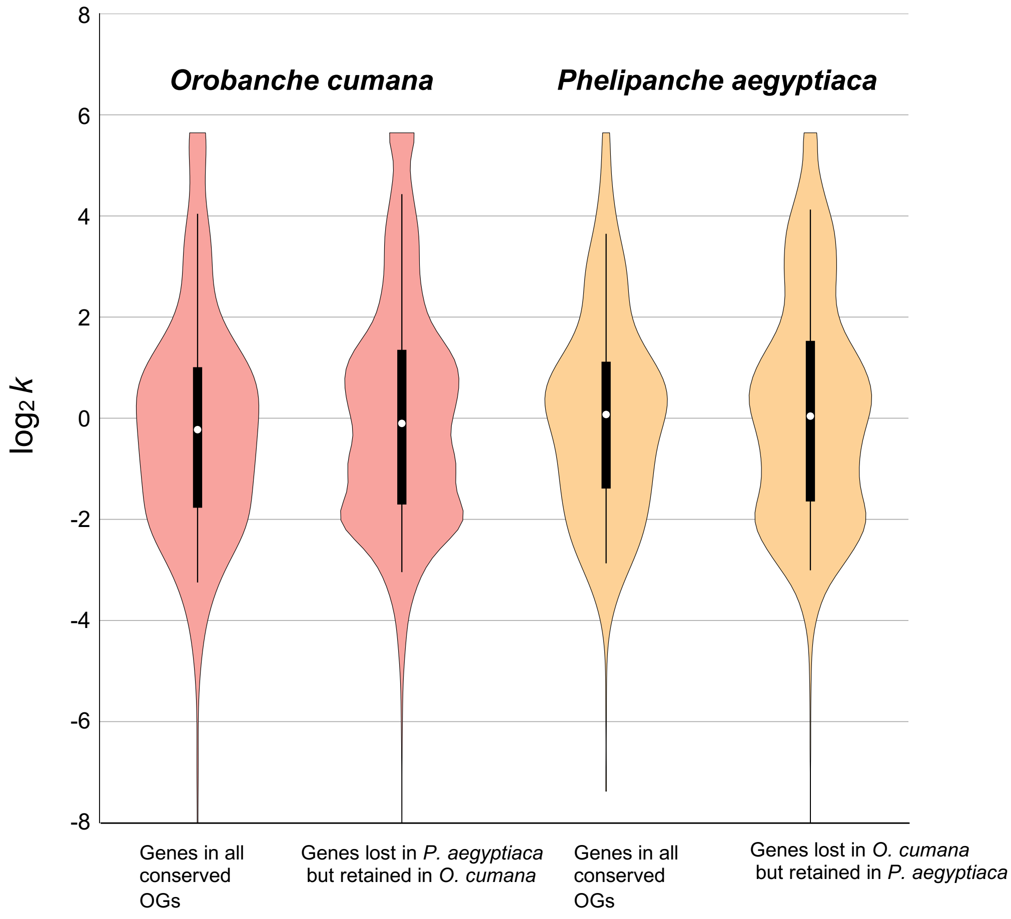


**Figure S9.** *k* value distributions of different gene sets in *Orobanche cumana* and *Phelipanche aegyptiaca*. Parameter *k* was calculated based on the general descriptive RELAX model (Wertheim et al., 2015). $k < 1$ ($\log_2 k < 0$) suggests a relaxation of purifying selection, whereas $k > 1$ ($\log_2 k > 0$) suggests selection intensification.

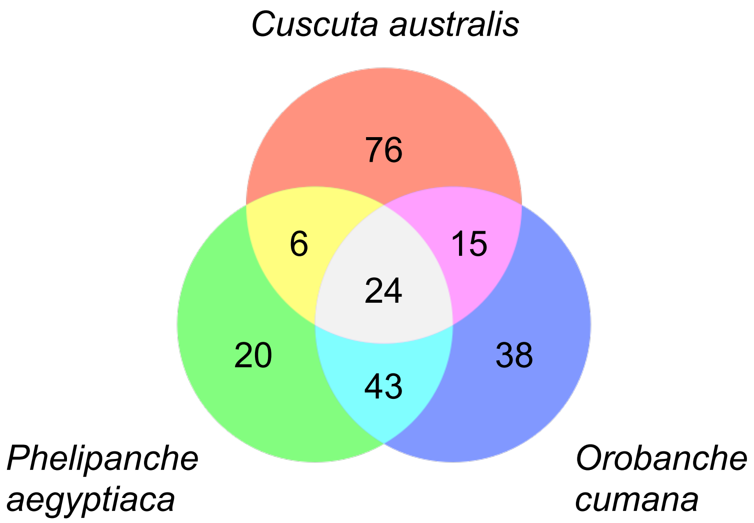


**Figure S10.** The Venn analysis of enriched Gene Ontology terms for lost genes in *Cuscuta australis*, *Phelipanche aegyptiaca* and *Orobanche cumana*.

**Table S1.** Statistics of survey, assembly, accuracy, and heterozygosity of *Lindenbergia luchunensis*, *Orobanche cumana*, and *Phelipanche aegyptiaca* genome

| | ***Lindenbergia luchunensis*** | ***Orobanche cumana*** | ***Phelipanche aegyptiaca*** |
|---|---|---|---|
| **Survey** | | | |
| Genome Size by flow cytometry | 223 Mb | 1,437.66 Mb | 3,992.35 Mb |
| Genome Size by *k*-mer | 244.44 Mb | 1,787.89 Mb | 3,956.65 Mb |
| Heterozygosity | 0.074% | 0.27% | 0.45% |
| Repeat Content | 44.6% | 84.54% | 84.76% |
| **Contig** | | | |
| short reads mapping ratio | 99.58% | 93.60% | 91.61% |
| Homozygosity (error) | 27623 bp | 173281 bp | 2709796 bp |
| | 0.01199% | 0.01214% | 0.07005% |
| N50 length | 8,232,122 bp | 13,334,487 bp | 9,973,504 bp |
| Total length | 243,957,151 bp | 1,462,975,560 bp | 3,876,900,029 bp |
| **Chromosome Level** | | | |
| Pseudo-chromosome number | 20 | 19 | - |
| Unanchored length | 28,764,949 bp | 42,580,799 bp | - |
| Organelle sequence length | 1,192,137 bp | 2,654,956 bp | - |
| Pseudo-chromosome length | 212,211,736 bp | 1,417,936,538 bp | - |

**Table S2.** Statistics of repetitive element of *Lindenbergia luchunensis*, *Orobanche cumana*, and *Phelipanche aegyptiaca* genome

| | *Lindenbergia luchunensis* | | | *Orobanche cumana* | | | *Phelipanche aegyptiaca* | | |
|---|---|---|---|---|---|---|---|---|---|
| | Count | Length (bp) | ratio | Count | Length (bp) | ratio | Count | Length (bp) | ratio |
| **LTR** | | | | | | | | | |
| Copia | 16,367 | 13,501,938 | 5.58% | 357,859 | 297,629,899 | 20.36% | 1,114,182 | 1,319,485,322 | 34.03% |
| Gypsy | 6,255 | 5,496,977 | 2.27% | 338,334 | 348,077,032 | 23.81% | 868,059 | 759,089,712 | 19.58% |
| unknown | 9,065 | 3,768,705 | 1.56% | 462,622 | 270,558,058 | 18.51% | 918,261 | 471,368,305 | 12.16% |
| **TIR** | | | | | | | | | |
| CACTA | 11,982 | 3,178,491 | 1.31% | 91,932 | 40,518,150 | 2.77% | 382,716 | 198,419,553 | 5.12% |
| Mutator | 39,811 | 13,875,978 | 5.74% | 83,998 | 31,927,033 | 2.18% | 312,101 | 133,186,489 | 3.44% |
| PIF_Harbinger | 1,800 | 526,169 | 0.22% | 15,931 | 5,412,412 | 0.37% | 85,001 | 35,186,043 | 0.91% |
| Tc1_Mariner | 1,855 | 397,540 | 0.16% | 70,604 | 18,803,223 | 1.29% | 62,323 | 25,011,704 | 0.65% |
| hAT | 19,882 | 5,849,970 | 2.42% | 71,575 | 63,160,203 | 4.32% | 79,103 | 33,731,360 | 0.87% |
| **nonLTR** | | | | | | | | | |
| LINE_element | 684 | 536,730 | 0.22% | 8,586 | 5,030,467 | 0.34% | 6,095 | 2,454,116 | 0.06% |
| unknown | 209 | 356,756 | 0.15% | 0 | 0 | 0.00% | 111 | 103,804 | 0.00% |
| **nonTIR** | | | | | | | | | |
| helitron | 75,133 | 23,475,291 | 9.70% | 125,462 | 52,810,974 | 3.61% | 395,347 | 191,055,088 | 4.93% |
| **Repeat_region** | 10,770 | 3,342,147 | 1.38% | 104,060 | 33,057,492 | 2.26% | 356,145 | 103,122,466 | 2.66% |
| **Total** | 193,813 | 74,306,692 | 30.72% | 1,730,963 | 1,166,984,943 | 79.82% | 4,579,444 | 3,272,213,961 | 84.40% |

**Table S3.** RNA-seq data used for genome annotation

| Sample ID | Tissue Description | Sample Source Species | Annotated Genome | Data Source |
|---|---|---|---|---|
| Ye03 | Seedling | *Lindenbergia muraria* | *Lindenbergia luchunensis* | This work |
| Ye07 | Root | *Lindenbergia muraria* | *Lindenbergia luchunensis* | This work |
| Ye16 | Leaf | *Lindenbergia muraria* | *Lindenbergia luchunensis* | This work |
| Ye18 | Flower | *Lindenbergia muraria* | *Lindenbergia luchunensis* | This work |
| Ye23 | Bud | *Lindenbergia muraria* | *Lindenbergia luchunensis* | This work |
| Ye73 | Seed | *Lindenbergia muraria* | *Lindenbergia luchunensis* | This work |
| YYe09 | Stem | *Lindenbergia muraria* | *Lindenbergia luchunensis* | This work |
| LC | Leaf, bud, and stem | *Lindenbergia luchunensis* | *Lindenbergia luchunensis* | This work |
| ZJX5232 | Pre-haustoria | *Orobanche cumana* | *Orobanche cumana* | This work |
| ZJX5235 | Seed | *Orobanche cumana* | *Orobanche cumana* | This work |
| ZJX5236 | Haustoria | *Orobanche cumana* | *Orobanche cumana* | This work |
| ZJX5237 | Seeding | *Orobanche cumana* | *Orobanche cumana* | This work |
| ZJX5238 | Stem | *Orobanche cumana* | *Orobanche cumana* | This work |
| ZJX5239 | Bud | *Orobanche cumana* | *Orobanche cumana* | This work |
| ZJX5240 | Flower | *Orobanche cumana* | *Orobanche cumana* | This work |
| ZJX5241 | Fruit | *Orobanche cumana* | *Orobanche cumana* | This work |
| PhAe_0 | Seed germination | *Phelipanche aegyptiaca* | *Phelipanche aegyptiaca* | PPGP |
| PhAe_1 | Root pre-haustorial initiation | *Phelipanche aegyptiaca* | *Phelipanche aegyptiaca* | PPGP |
| PhAe_2 | Root post-haustorial initiation | *Phelipanche aegyptiaca* | *Phelipanche aegyptiaca* | PPGP |
| PhAe_3 | Haustoria attached to host | *Phelipanche aegyptiaca* | *Phelipanche aegyptiaca* | PPGP |
| PhAe_4_1 | Vascular early development | *Phelipanche aegyptiaca* | *Phelipanche aegyptiaca* | PPGP |
| PhAe_4_2 | Vascular late development | *Phelipanche aegyptiaca* | *Phelipanche aegyptiaca* | PPGP |
| PhAe_5_1 | Pre-emerged leaf and stem | *Phelipanche aegyptiaca* | *Phelipanche aegyptiaca* | PPGP |
| PhAe_5_2 | Root | *Phelipanche aegyptiaca* | *Phelipanche aegyptiaca* | PPGP |
| PhAe_6_1 | Emerged leaf and stem | *Phelipanche aegyptiaca* | *Phelipanche aegyptiaca* | PPGP |
| PhAe_6_2 | Reproductive structure | *Phelipanche aegyptiaca* | *Phelipanche aegyptiaca* | PPGP |

* PPGP (http://ppgp.huck.psu.edu)

**Table S4.** Results of BUSCO analysis for the genome and protein-coding gene set of *Lindenbergia luchunensis*, *Orobanche cumana*, and *Phelipanche aegyptiaca* genome

| | Complete BUSCOs | | | Fragmented BUSCOs | Missing BUSCOs | Total BUSCO Groups Searched |
|---|---|---|---|---|---|---|
| | Complete | Single copy | Duplicated | | | |
| ***Lindenbergia luchunensis*** | 97.1% | 91.4% | 5.7% | 1.0% | 1.9% | 1614 |
| ***Orobanche cumana*** | 80.5% | 77.6% | 2.9% | 1.8% | 17.7% | 1614 |
| ***Phelipanche aegyptiaca*** | 76.3% | 65.4% | 10.9% | 3.3% | 20.4% | 1614 |

**Table S5.** Plant genomes and transcriptomes for the phylogenetic analysis

| Species Name | Abbr. | Order Name | Lifestyle | Data Source (NCBI Bioproject) | Reference |
|---|---|---|---|---|---|
| *Coffea canephora* | Cca | Outgroup | autotrophic | PRJEB4211 | (Denoeud et al., 2014) |
| *Olea europaea* | Oeu | Outgroup | autotrophic | PRJNA417827 | (Rao et al., 2021) |
| *Mimulus guttatus* | Mgu | Outgroup | autotrophic | PRJNA13880 | (Hellsten et al., 2013) |
| *Rehmannia glutinosa* | Rgl | Rehmannieae | autotrophic | PRJEB21674 | (Ma et al., 2021) |
| *Lindenbergia luchunensis* | Llu | Lindenbergieae | autotrophic | This work | |
| *Monochasma savatieri* | Msa | Cymbarieae | hemiparasitic, facultative | PRJNA600790 | (Chen et al., 2021) |
| *Cymbaria mongolica* | Cmo | Cymbarieae | hemiparasitic, facultative | PRJNA636634 | (Zhang et al., 2020) |
| *Melampyrum roseum* | Mro | Rhinantheae | hemiparasitic, facultative | PRJDB5395 | (Kado and Innan, 2018) |
| *Euphrasia arctica* | Ear | Rhinantheae | hemiparasitic, facultative | PRJNA624746 | (Becher et al., 2020) |
| *Bartsia alpine* | Bal | Rhinantheae | hemiparasitic, facultative | PRJNA622490 | (Marx et al., 2021) |
| *Pedicularis cranolopha* | Pcr | Pediculardeae | hemiparasitic, facultative | PRJNA815880 | |
| *Phtheirospermum japonicum* | Pja | Pedicularideae | hemiparasitic, facultative | PRJDB10085 | (Cui et al., 2020) |
| *Aureolaria pectinate* | Ape | Pedicularideae | hemiparasitic, facultative | PRJNA359989 | (Boachon et al., 2018) |
| *Triphysaria versicolor* | Tve | Pedicularideae | hemiparasitic, facultative | PRJNA340868 | (Westwood et al., 2012) |

| | | | | | |
|---|---|---|---|---|---|
| *Striga asiatica* | Sas | Buchnereae | hemiparasitic, obligate | PRJDB4996 | (Yoshida et al., 2019) |
| *Aeginetia indica* | Ain | Buchnereae | holoparasitic, obligate | PRJNA556535 | (Chen et al., 2020) |
| *Alectra vogelii* | Avo | Buchnereae | hemiparasitic, obligate | PRJNA340868 | (Wickett et al., 2011) |
| *Centranthera grandiflora* | Cgr | Buchnereae | hemiparasitic, facultative | PRJNA558809 | (Zhang et al., 2019) |
| *Cistanche deserticola* | Cde | Orobancheae | holoparasitic, obligate | PRJNA598927 | |
| *Orobanche cumana* | Ocu | Orobancheae | holoparasitic, obligate | This work | |
| *Aphyllon californicum* | Aca | Orobancheae | holoparasitic, obligate | PRJNA340868 | (Yang et al., 2015) |
| *Phelipanche aegyptiaca* | Pae | Orobancheae | holoparasitic, obligate | This work | |

**Table S6.** Statistics of gene duplication types in *Pedicularis cranolopha*, *Phtheirospermum japonicum, Striga asiatica, Orobanche cumana*, and *Phelipanche aegyptiaca*.

| Species Name | Total Gene Number | Number of Orthogroups Used for Statistics | Segmental Duplication | Tandem Duplication | LTR-assisted Duplication |
|---|---|---|---|---|---|
| *Pedicularis cranolopha* | 47689 | 1500 | 209 (13.93%) | 636 (42.40%) | 165 (11.00%) |
| *Phtheirospermum japonicum* | 30264 | 1006 | 38 (3.78%) | 302 (30.02%) | 175 (17.40%) |
| *Striga asiatica* | 33426 | 1997 | 1129 (56.53%) | 180 (9.01%) | 120 (6.01%) |
| *Orobanche cumana* | 42525 | 493 | 36 (7.30%) | 176 (35.70%) | 81 (16.43%) |
| *Phelipanche aegyptiaca* | 50484 | 1634 | 282 (17.26%) | 224 (13.71%) | 107 (6.55%) |

Note: For the selection criteria of the OGs involved in the statistics and the identification of various types of gene duplication, please refer to Supplemental Methods.

**Table S7.** RNA-seq data used for identification of haustorially highly expressed genes.

| Species | Sample ID | Tissue Description | Group | Data Source | Reference |
|---|---|---|---|---|---|
| *Striga asiatica* | StHe_0_R1 | Seed germination | Non-haustorium group | PPGP | |
| *Striga asiatica* | StHe_0_R2 | Seed germination | Non-haustorium group | PPGP | |
| *Striga asiatica* | StHe_0_R3 | Seed germination | Non-haustorium group | PPGP | |
| *Striga asiatica* | StHe_1_R1 | Root pre-haustorial initiation | Non-haustorium group | PPGP | |
| *Striga asiatica* | StHe_1_R2 | Root pre-haustorial initiation | Non-haustorium group | PPGP | |
| *Striga asiatica* | StHe_1_R3 | Root pre-haustorial initiation | Non-haustorium group | PPGP | |
| *Striga asiatica* | StHe_2_R1 | Root post-haustorial initiation | Haustorium group | PPGP | |
| *Striga asiatica* | StHe_2_R2 | Root post-haustorial initiation | Haustorium group | PPGP | |
| *Striga asiatica* | StHe_2_R3 | Root post-haustorial initiation | Haustorium group | PPGP | |
| *Striga asiatica* | StHe_3_R1 | Haustoria attached to host root | Haustorium group | PPGP | |
| *Striga asiatica* | StHe_3_R2 | Haustoria attached to host root | Haustorium group | PPGP | |
| *Striga asiatica* | StHe_3_R3 | Haustoria attached to host root | Haustorium group | PPGP | (Westwood et al., 2012) |
| *Striga asiatica* | StHe_4_R1 | Vascular late development | Haustorium group | PPGP | |
| *Striga asiatica* | StHe_4_R2 | Vascular late development | Haustorium group | PPGP | |
| *Striga asiatica* | StHe_4_R3 | Vascular late development | Haustorium group | PPGP | |
| *Striga asiatica* | StHe_5_R1 | Roots | Non-haustorium group | PPGP | |
| *Striga asiatica* | StHe_5_R2 | Roots | Non-haustorium group | PPGP | |
| *Striga asiatica* | StHe_5_R3 | Roots | Non-haustorium group | PPGP | |
| *Striga asiatica* | StHe_6_1_R1 | Emerged leaves and stems | Non-haustorium group | PPGP | |
| *Striga asiatica* | StHe_6_1_R2 | Emerged leaves and stems | Non-haustorium group | PPGP | |
| *Striga asiatica* | StHe_6_1_R3 | Emerged leaves and stems | Non-haustorium group | PPGP | |
| *Striga asiatica* | StHe_6_2_R1 | Reproductive structures | Non-haustorium group | PPGP | |
| *Striga asiatica* | StHe_6_2_R2 | Reproductive structures | Non-haustorium group | PPGP | |
| *Striga asiatica* | StHe_6_2_R3 | Reproductive structures | Non-haustorium group | PPGP | |

| | | | | | |
|---|---|---|---|---|---|
| *Phelipanche aegyptiaca* | PhAe_0_R2 | Seed germination | Non-haustorium group | PPGP | |
| *Phelipanche aegyptiaca* | PhAe_0_R3 | Seed germination | Non-haustorium group | PPGP | |
| *Phelipanche aegyptiaca* | PhAe_0_R5 | Seed germination | Non-haustorium group | PPGP | |
| *Phelipanche aegyptiaca* | PhAe_1_R2 | Root pre-haustorial initiation | Non-haustorium group | PPGP | |
| *Phelipanche aegyptiaca* | PhAe_1_R3 | Root pre-haustorial initiation | Non-haustorium group | PPGP | |
| *Phelipanche aegyptiaca* | PhAe_1_R5 | Root pre-haustorial initiation | Non-haustorium group | PPGP | |
| *Phelipanche aegyptiaca* | PhAe_2_R3 | Root post-haustorial initiation | Haustorium group | PPGP | |
| *Phelipanche aegyptiaca* | PhAe_2_R4 | Root post-haustorial initiation | Haustorium group | PPGP | |
| *Phelipanche aegyptiaca* | PhAe_2_R5 | Root post-haustorial initiation | Haustorium group | PPGP | |
| *Phelipanche aegyptiaca* | PhAe_3_R2 | Haustoria attached to host | Haustorium group | PPGP | |
| *Phelipanche aegyptiaca* | PhAe_3_R3 | Haustoria attached to host | Haustorium group | PPGP | |
| *Phelipanche aegyptiaca* | PhAe_3_R4 | Haustoria attached to host | Haustorium group | PPGP | |
| *Phelipanche aegyptiaca* | PhAe_4_1_R2 | Vascular early development | Haustorium group | PPGP | (Westwood et al., 2012) |
| *Phelipanche aegyptiaca* | PhAe_4_1_R3 | Vascular early development | Haustorium group | PPGP | |
| *Phelipanche aegyptiaca* | PhAe_4_1_R4 | Vascular early development | Haustorium group | PPGP | |
| *Phelipanche aegyptiaca* | PhAe_4_2_R1 | Vascular late development | Haustorium group | PPGP | |
| *Phelipanche aegyptiaca* | PhAe_4_2_R3 | Vascular late development | Haustorium group | PPGP | |
| *Phelipanche aegyptiaca* | PhAe_4_2_R4 | Vascular late development | Haustorium group | PPGP | |
| *Phelipanche aegyptiaca* | PhAe_5_1_R1 | Pre-emerged leaves and stems | Non-haustorium group | PPGP | |
| *Phelipanche aegyptiaca* | PhAe_5_1_R3 | Pre-emerged leaves and stems | Non-haustorium group | PPGP | |
| *Phelipanche aegyptiaca* | PhAe_5_1_R5 | Pre-emerged leaves and stems | Non-haustorium group | PPGP | |
| *Phelipanche aegyptiaca* | PhAe_5_2_R1 | Roots | Non-haustorium group | PPGP | |
| *Phelipanche aegyptiaca* | PhAe_5_2_R2 | Roots | Non-haustorium group | PPGP | |
| *Phelipanche aegyptiaca* | PhAe_5_2_R5 | Roots | Non-haustorium group | PPGP | |
| *Phelipanche aegyptiaca* | PhAe_6_1_R3 | Emerged leaves and stems | Non-haustorium group | PPGP | |
| *Phelipanche aegyptiaca* | PhAe_6_1_R5 | Emerged leaves and stems | Non-haustorium group | PPGP | |

| | | | | | |
|---|---|---|---|---|---|
| *Phelipanche aegyptiaca* | PhAe_6_1_R6 | Emerged leaves and stems | Non-haustorium group | PPGP | |
| *Phelipanche aegyptiaca* | PhAe_6_2_R1 | Reproductive structures | Non-haustorium group | PPGP | |
| *Phelipanche aegyptiaca* | PhAe_6_2_R2 | Reproductive structures | Non-haustorium group | PPGP | |
| *Phelipanche aegyptiaca* | PhAe_6_2_R3 | Reproductive structures | Non-haustorium group | PPGP | |
| *Phtheirospermum japonicum* | DRR234959 | Roots | Non-haustorium group | NCBI | |
| *Phtheirospermum japonicum* | DRR234960 | Roots | Non-haustorium group | NCBI | |
| *Phtheirospermum japonicum* | DRR234961 | Roots | Non-haustorium group | NCBI | |
| *Phtheirospermum japonicum* | DRR234965 | 1-day post infection roots | Haustorium group | NCBI | |
| *Phtheirospermum japonicum* | DRR234966 | 1-day post infection roots | Haustorium group | NCBI | (Cui et al., 2020) |
| *Phtheirospermum japonicum* | DRR234967 | 1-day post infection roots | Haustorium group | NCBI | |
| *Phtheirospermum japonicum* | DRR234968 | 7-day post infection roots | Haustorium group | NCBI | |
| *Phtheirospermum japonicum* | DRR234969 | 7-day post infection roots | Haustorium group | NCBI | |
| *Phtheirospermum japonicum* | DRR234970 | 7-day post infection roots | Haustorium group | NCBI | |

**Table S8.** Haustorially highly expressed genes in *Phtheirospermum japonicum*, *Striga asiatica*, and *Phelipanche aegyptiaca*.

| | ***Phtheirospermum japonicum*** | ***Striga asiatica*** | ***Phelipanche aegyptiaca*** |
|---|---|---|---|
| Duplicated after MRCA of Orobanchaceae and *Coffea* | 310 | 629 | 498 |
| Non-duplicated after MRCA of Orobanchaceae and *Coffea* | 418 | 908 | 684 |
| Total highly expressed genes in haustoria | 728 | 1537 | 1182 |

**Table S9.** Phylogenetic placement of gene duplications detected in the haustorially highly expressed genes.

* Numbers represent the number of orthogroups with duplication events.

| **Phylogenetic placement** | ***Phtheirospermum japonicum*** | ***Striga asiatica*** | ***Phelipanche aegyptiaca*** |
|---|---|---|---|
| species-specific | 81 | 306 | 258 |
| MRCA of Orobancheae | - | - | 16 |
| MRCA of Orobancheae and *Striga asiatica* | - | 1 | 4 |
| MRCA of Pedicularideae | 30 | - | - |
| MRCA of orobanchaceous parasites | 29 | 23 | 14 |
| MRCA of Orobanchaceae | 31 | 42 | 42 |
| MRCA of Orobanchaceae and *Mimulus guttatus* ($\beta^{L}$ WGD event) | 121 | 184 | 148 |
| MRCA of Lamiales and *Solanum lycopersicum* | 4 | 9 | 8 |
| MRCA of asterids | 37 | 64 | 38 |
| Total duplicated | 333 | 629 | 528 |

**Table S10.** Survival of gene duplicates caused by WGD events in different species.

* see Supplemental Methods for details about Exponential Model.

| Species | Scope of Statistics | Orthogroups with retained gene duplicates | Orthogroups | Percentage of retained gene duplicates |
|---|---|---|---|---|
| *Mimulus guttatus* | Genome-wide | 2333 | 14658 | 15.92% |
| *Lindenbergia luchunensis* | Genome-wide | 2498 | 13856 | 18.03% |
| *Phtheirospermum japonicum* | Genome-wide | 1942 | 13089 | 14.84% |
| | Haustoria-related orthogroups | 121 | 401 | 30.17% |
| *Striga asiatica* | Genome-wide | 1138 | 12080 | 9.42% |
| | Haustoria-related orthogroups | 184 | 1024 | 17.97% |
| *Phelipanche aegyptiaca* | Genome-wide | 1033 | 11405 | 9.06% |
| | Haustoria-related orthogroups | 148 | 844 | 17.54% |
| Exponential Model * | Today | Ks = 0.835 ($\beta^{L}$); *a* = 0.13020; *b* = 0 | | 18.23% |
| | MRCA of orobanchaceous parasites | Ks = 0.835 ($\beta^{L}$) - 0.476 (MRCA of orobanchaceous parasites); *a* = 0.13020; *b* = 0 | | 48.10% |

**Dataset S1 (separate file).** Gene family expansions and contractions

**Dataset S2 (separate file).** Results of GO enrichment analyses

Dataset S2a. GO enrichment of the orthogroups which experienced expansions when parasitism first appeared in Orobanchaceae

Dataset S2b. GO enrichment of the orthogroups which experienced expansions when obligate parasitism first appeared in Orobanchaceae

Dataset S2c. GO enrichment of the orthogroups which experienced expansions when holoparasitism first appeared in Orobanchaceae

Dataset S2d. GO enrichment of the orthogroups which experienced expansions in *Phtheirospermum japonicum*

Dataset S2e. GO enrichment of the orthogroups which experienced expansions in *Pedicularis cranolopha*

Dataset S2f. GO enrichment of the orthogroups which experienced expansions in *Striga asiatica*

Dataset S2g. GO enrichment of the orthogroups which experienced expansions in *Orobanche cumana*

Dataset S2h. GO enrichment of the orthogroups which experienced expansions in *Phelipanche aegyptiaca*

**Dataset S3 (separate file).** Gene loss identified using Arabidopsis genes as reference
